# Human Replication Protein A complex is a Telomerase Processivity Factor Essential for Telomere Maintenance

**DOI:** 10.1101/2024.08.16.608355

**Authors:** Sourav Agrawal, Xiuhua Lin, Vivek Susvirkar, Michael S. O’Connor, Bianca L. Chavez, Victoria R. Tholkes, Kaitlyn M. Abe, Qixiang He, Xuhui Huang, Ci Ji Lim

## Abstract

Telomerase is crucial for maintaining telomere length and safeguarding genome stability. In this study, we identified Replication Protein A (RPA) as a novel telomerase processivity factor, functioning alongside the telomerase recruitment factor TPP1-POT1. AlphaFold2 predictions revealed that RPA and TPP1 interact with telomerase at distinct binding sites. Using separation-of-function mutants, we discovered that RPA-mediated telomerase stimulation is indispensable for telomere elongation, while TPP1-POT1 primarily functions in recruiting telomerase to telomeres. Furthermore, we demonstrated that short telomere disease-associated telomerase mutations compromise RPA’s ability to stimulate telomerase, establishing a link between impaired RPA-dependent processivity and telomeropathies. Our findings redefine human telomerase regulation by establishing RPA as a critical regulator and provide new insights into the molecular basis of telomere-related diseases.

---

Telomeres, the natural ends of linear chromosomes, are crucial for maintaining genome stability (*1*). In humans, telomere maintenance (2, 3) involves the extension of the G-rich single-stranded DNA (4) (ssDNA) by telomerase (5–8) and the synthesis of the complementary C-strand by DNA polymerase α-primase (Polα-primase) in conjunction with the CST complex (9, 10). Telomerase is recruited to telomeres by the shelterin protein TPP1 (11–16), which together with POT1 also stimulates telomerase repeat addition processivity (RAP) in vitro (17, 18). However, the role of this stimulatory function in telomere maintenance in vivo remains unclear.

Replication protein A (RPA) is a highly conserved heterotrimeric complex involved in DNA replication and repair (19, 20). In ciliates and yeast, RPA-like complexes are implicated in telomerase regulation and telomere maintenance (21–27). The human CST complex, despite structural similarities to RPA (28–30), is known to terminate rather than promote telomerase activity (31, 32). Given these observations, we investigated whether human RPA directly regulates telomerase activity.

## RPA stimulates telomerase processivity through ssDNA and TERT interactions

Using primer-extension assays, we found that recombinant human RPA (**fig. S1**) increased telomerase repeat addition processivity (RAP) by approximately six-fold, extending primers to ∼130 nucleotides (nt), similar to the effect of TIN2-TPP1-POT1 (TPT) addition (**Fig. 1A & 1B**). Like TPT addition, RPA addition did not affect overall telomerase activity (**Fig. 1C**)(17), indicating RPA’s effect on telomerase enzymatic activity is restricted to RAP stimulation.

**Fig. 1.**
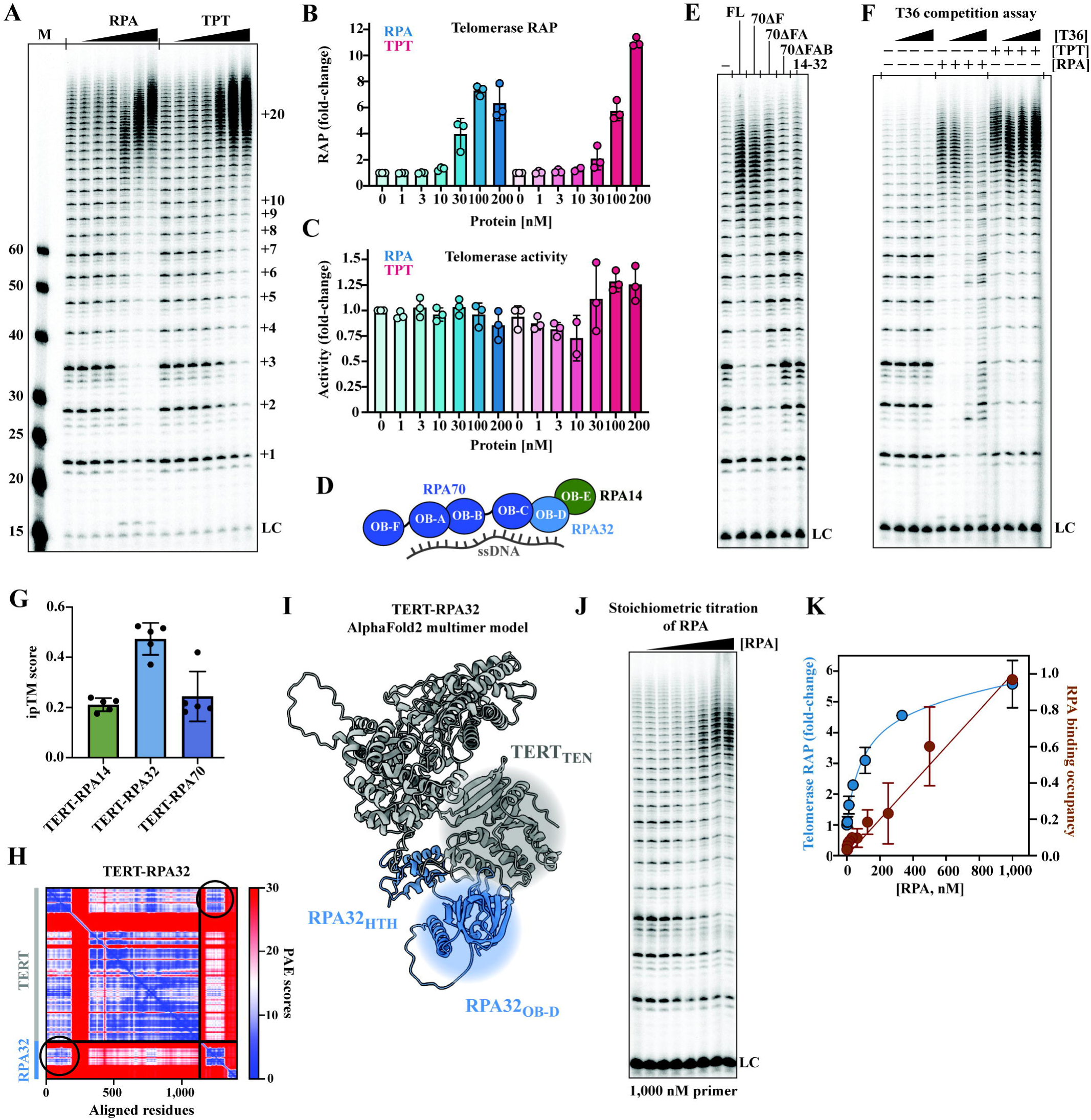
Human RPA stimulates telomerase processivity through ssDNA and TERT interactions. **(A)** Direct telomerase assay comparing the effects of increasing concentrations of RPA and TPT (TPP1-POT1-TIN2). Denaturing gel electrophoresis analysis shows telomere repeat addition products with ladder-like banding patterns. Numbers on the right indicate the number of telomere repeats added. **(B)** Quantification of telomerase repeat addition processivity (RAP) in response to RPA (blue) and TPT (pink) concentrations ranging from 0 to 200 nM. RAP is expressed as fold-change and relative to a no added protein control. **(C)** Quantification of total telomerase activity in response to RPA and TPT concentrations (0-200 nM), shown as fold- change in activity. The fold-change is expressed relative to the telomerase alone lane. **(D)** Schematic representation of a human RPA heterotrimer binding to a single-stranded DNA (ssDNA), showing the arrangement of OB-fold domains in RPA70, RPA32, and RPA14 subunits. **(E)** Direct telomerase assays using various RPA constructs: full-length (FL), RPA with RPA70ΔF, RPA70ΔFA, and RPA70ΔFAB, as well as the RPA14-32 heterodimer. **(F)** Competition assay showing the effects of increasing concentrations of dT_36_ DNA oligo (T36) competitor on telomerase activity in the presence of RPA and TPT. **(G)** AlphaFold2 predictions integrated predicted TM-score (ipTM) for TERT interactions with different RPA subunits (RPA14, RPA32, RPA70), indicating confidence in predicted structural models. **(H)** Predicted Aligned Error (PAE) plot for the TERT-RPA32 interaction, showing the expected distance error (Ångstroms) between residue pairs. Lower values (blue) indicate higher confidence. **(I)** AlphaFold2 multimer model of the TERT-RPA32 complex, highlighting the predicted structural arrangement of TERT (gray) with RPA32 OB-D domain (blue). **(J)** Stoichiometric titration of RPA in direct telomerase assay, showing gel electrophoresis results with increasing RPA concentrations up to 1,000 nM. Primer concentration is 1,000 nM. **(K)** Quantification of telomerase RAP (blue, left y-axis) and RPA binding occupancy (red, right y-axis) in response to increasing RPA concentrations (0-1,000 nM). LC denotes loading control. Error bars represent standard deviation from multiple experiments. M in panel **A** indicates molecular weight markers.

To determine if the full RPA heterotrimeric complex (RPA70-RPA32-RPA14) (**Fig. 1D**) is required for RAP stimulation, we tested individual subunits and subcomplexes (**fig. S2A**). Neither RPA70 alone, the RPA14-32 subcomplex, nor the RPA trimeric core (33) significantly affected telomerase activity or processivity (**fig. S2B-D**). These results demonstrate that an intact RPA heterotrimer is required to function as a telomerase processivity factor.

To understand how RPA stimulates telomerase RAP, we first investigated the role of RPA’s ssDNA-binding domains. We generated RPA complexes with truncated RPA70 OB domains (**Fig. 1D & fig. S2A**) and tested their ability to stimulate telomerase RAP. Removal of the RPA70 OB-F domain, which has minimal ssDNA-binding affinity (34), did not affect RAP stimulation (**Fig. 1E**). However, deletion of the OB-A and OB-B domains, crucial for RPA ssDNA binding (35, 36), reduced or abolished RPA’s stimulatory effect (**Fig. 1E**). These findings suggest RPA ssDNA binding is critical for its stimulation of telomerase RAP.

To further test the importance of ssDNA binding (37), we designed a competitive inhibition experiment using a polyT oligonucleotide (T36), leveraging on RPA’s ssDNA binding sequence promiscuity (37). We reasoned that if RPA’s ssDNA binding is crucial for its telomerase RAP stimulation, adding the T36 oligos will sequester RPA from binding to the telomeric primer, leading to a loss in telomerase RAP stimulation. Indeed, we saw T36 oligo titration led to a reduction in RPA stimulation of telomerase RAP (**Fig. 1F**), furthering underlining the importance of RPA ssDNA binding for its telomerase RAP stimulation. Control experiments showed T36 oligo titration did not affect telomerase activity or TPT stimulation (**Fig. 1F**), as expected given telomerase and POT1’s binding preference for telomeric sequences (38, 39).

TPP1-POT1 requires both protein-protein and protein-DNA interactions for telomerase RAP stimulation (18). We therefore examined if RPA also has direct interactions with TERT. To investigate this, we used AlphaFold2 (40) to screen for potential protein-protein interactions between RPA subunits and TERT. The software predicted that the RPA32 OB-D domain interacts with TERT TEN domain(41) (**Fig. 1G-I**).

To test this prediction, we performed stoichiometric titrations of RPA to the telomerase DNA primer in a telomerase assay. If there is additional RPA-TERT interaction beyond ssDNA-binding as predicted by AlphaFold2, we should observe a non-linear increase in RPA telomerase RAP stimulation with stoichiometric titration of RPA. Conversely, a purely ssDNA binding-dependent mechanism would yield a linear increase in telomerase RAP stimulation with increasing RPA concentration.

We observed a non-linear response (**Fig. 1J & K**), with almost peak RAP stimulation at substoichiometric RPA binding concentrations. This non-linear response resembles that seen with TPP1-POT1 (18) and supports our AlphaFold2 prediction that RPA directly interacts with TERT. A stoichiometric binding assay using similar conditions showed a linear response in RPA binding to the telomeric DNA primer (**Fig. 1K**), indicating that the substoichiometric stimulation is not due to poor RPA binding or protein concentration miscalculation.

Together, these results demonstrate that RPA stimulates telomerase RAP through both ssDNA binding and protein-protein interactions, similar to the mechanism of TPP1-POT1.

## RPA32 OB-D and TPP1 OB domains bind telomerase at separate sites

To determine whether RPA and TPP1-POT1 compete for the same telomerase binding site, we docked our RPA32-TERT AlphaFold2 model into cryo-EM structures of human telomerase bound to the TPP1 OB domain (41, 42). The analysis revealed that RPA32 OB-D and TPP1 OB domains bind to separate surfaces of TERT without clashes (**Fig. 2A**). RPA32 OB binds solely to the TERT TEN domain, while TPP1 OB domain spans the TEN and IFD domains (41–43). This binding arrangement is also predicted by AlphaFold2 when we included TPP1 OB domain in the prediction (**fig. S3**).

**Fig. 2.**
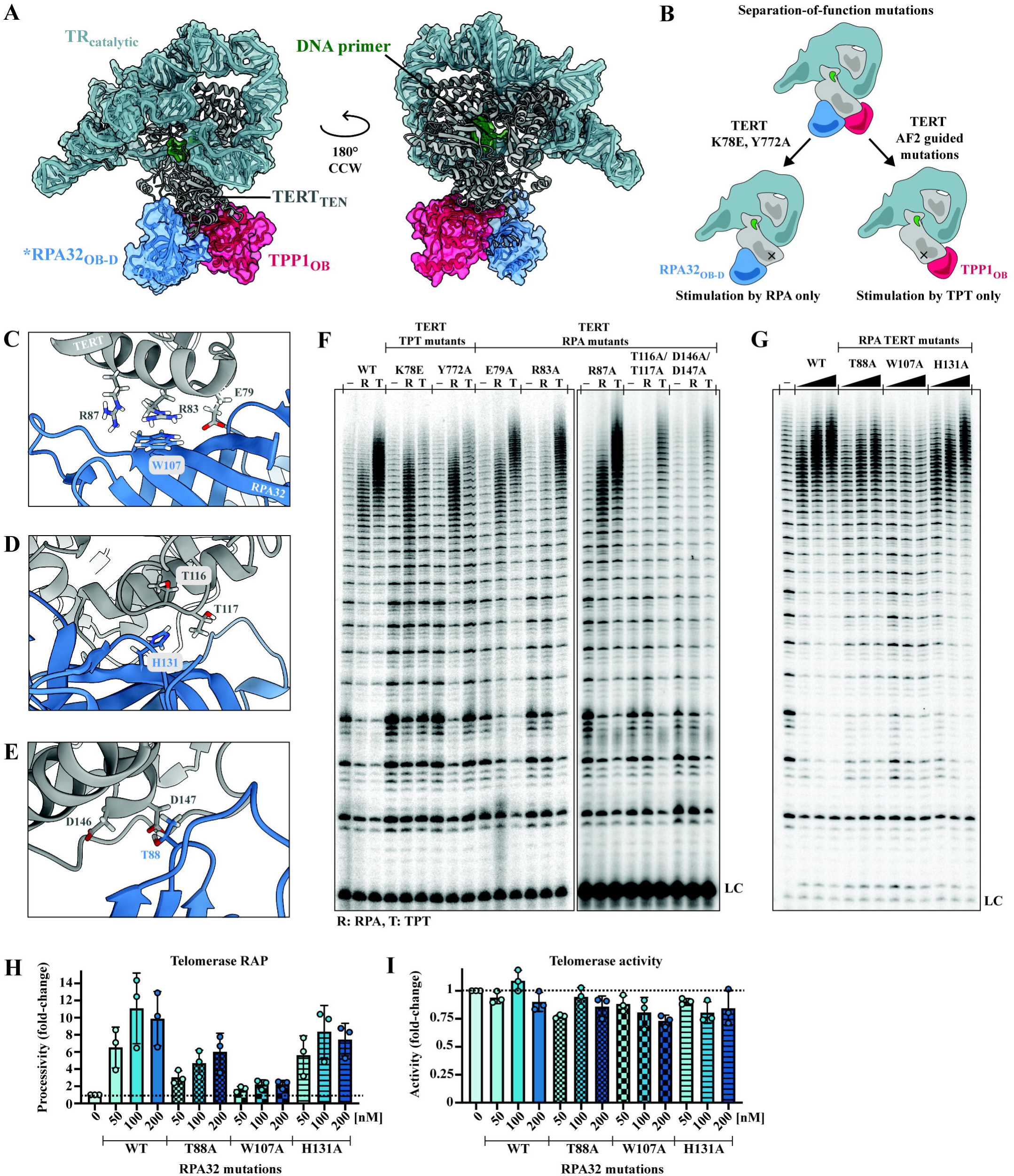
Distinct TERT binding sites for RPA32 OB-D and TPP1 OB domains enable separation-of-function mutations. **(A)** AlphaFold2 multimer model of the telomerase complex showing TERT (gray), TR_catalytic_ (light blue), DNA primer (green), RPA32_OB-D_ (blue), and TPP1_OB_ (red). Two views rotated 180° counterclockwise (CCW) are shown. Asterisk indicate AlphaFold2 predicted model. **(B)** Schematic representation of separation-of-function mutations in TERT affecting RPA or TPT stimulation. **(C-E)** Close-up views of predicted key interaction interfaces between TERT (gray) and RPA32 (blue), highlighting specific residues. **(F)** Direct telomerase assays comparing wild-type (WT) TERT with various TPT-interaction mutants (K78E, Y772A) and RPA-interaction mutants (E79A, R83A, R87A, T116A/D117A, D146A/D147A). **(G)** Direct telomerase assays with WT TERT and RPA complexes containing mutations in the RPA32 subunit (T88A, W107A, H131A). **(H)** Quantification of telomerase repeat addition processivity (RAP) for WT and RPA complexes with RPA32 mutations at varying concentrations (0-200 nM) from panel **G**. **(I)** Quantification of total telomerase activity for WT and RPA complexes with RPA32 mutations at varying concentrations (0-200 nM) from panel **G**. R: RPA, T: TPT. LC denotes loading control. Error bars represent standard deviation from multiple experiments.

The predicted distinct binding sites of RPA32 and TPP1 present us with an opportunity to identify separation-of-function (SOF) TERT mutants that would selectively disrupt RPA32 OB- D domain binding but not TPP1 OB domain binding, leading to selective telomerase RAP stimulation (**Fig. 2B**). To refine our predictions, we performed all-atom molecular dynamics simulations in explicit solvent on the AlphaFold2 model. This approach allowed us to relax the predicted protein-protein interactions and identify conserved TERT residues E79, R83, R87, T116, T117, D146, and D147 as potential RPA32 interaction sites. (**Fig. 2C-E, fig. S5A**).

We reconstituted telomerase complexes with TERT mutations E79A, R83A, R87A, T116A/T117A, and D146A/D147A and tested these mutants’ ability to be stimulated by RPA using a direct telomerase assay. We found that R83A, T116A/T117A, and D146A/D147A mutations selectively abolished RPA-mediated processivity stimulation without affecting TPT-mediated stimulation (**Fig. 2F**). All mutants had telomerase activity comparable to wild-type (WT) (normalized by telomerase RNA, TR, levels), so the observed effects were unlikely due to structural defects in the enzymes. These results thus identify R83A, T116A/T117A, and D146A/D147A as SOF mutants specific for disrupting RPA-mediated telomerase stimulation.

As a complementary approach, we also tested previously reported TERT TPT mutants K78E (44) and Y772A (42). Based on the AlphaFold2 prediction, disrupting the TERT-TPP1_OB_ interaction should not affect RPA-mediated telomerase RAP stimulation and only impair stimulation by TPT. Indeed, we found these mutations disrupted TPT-mediated stimulation and RPA-mediated telomerase RAP stimulation to a lesser extent, if not at all (**Fig. 2F**). The K78E suffered a reduction in RPA stimulation, possibly due to the proximity of the residue to the identified TERT-RPA32 interaction site (**Fig. 2C**). Y772A did not affect RPA telomerase RAP stimulation. These results further support the AlphaFold2 prediction that RPA32 OB-D and TPP1 OB domains bind to separate sites on TERT.

To further test the predicted TERT-RPA32 interactions, we identified RPA32 OB-D domain residues T88, W107, and H131 as potential interaction partners for TERT D146, R83, and T116, respectively (**Fig. 2C-E**). In particular, we saw that TERT R83 forms a pi-cation interaction with RPA32 W107 (**Fig. 2C**). We generated recombinant RPA complexes containing RPA32 T88A, W107A, and H131A mutations (**fig. S4 & S5B**) and tested their ability to stimulate telomerase RAP in a direct assay. We found the T88A and H131A mutants had a slightly lower telomerase RAP stimulatory effect than WT while the W107A mutant had a near-complete loss of telomerase RAP stimulation (**Fig. 2G-I**). The W107A mutation did not affect RPA’s ability to bind telomeric ssDNA (**fig. S6**) (45), indicating that the loss of RAP stimulation by the RPA32 W107A mutant is not due to defective ssDNA binding, but instead supports the AlphaFold2- predicted protein-protein interaction between TERT and RPA32 OB-D domain. The successful identification of these SOF mutants without an experimental structure underlines the power of AlphaFold2 in guiding functional studies of protein-protein interactions.

## RPA localizes at telomeres via telomerase without triggering a DNA damage response

To investigate the cellular function of RPA stimulation of telomerase RAP, we used immunofluorescence (IF) imaging to determine whether RPA localizes to telomeres upon telomerase recruitment. Our IF analysis revealed that endogenous RPA molecules (with RPA32 detection serving as a proxy) are indeed localized at telomeres in HeLa cell lines expressing telomerase (**Fig. 3A**). This telomeric localization of RPA required the expression of both TERT and telomerase RNA (TR), as TERT alone was insufficient to recruit RPA to telomeres (**Fig. 3A**). These results suggest that a fully assembled telomerase ribonucleoprotein complex is necessary either to recruit RPA to telomeres or to stabilize its localization there.

**Fig. 3.**
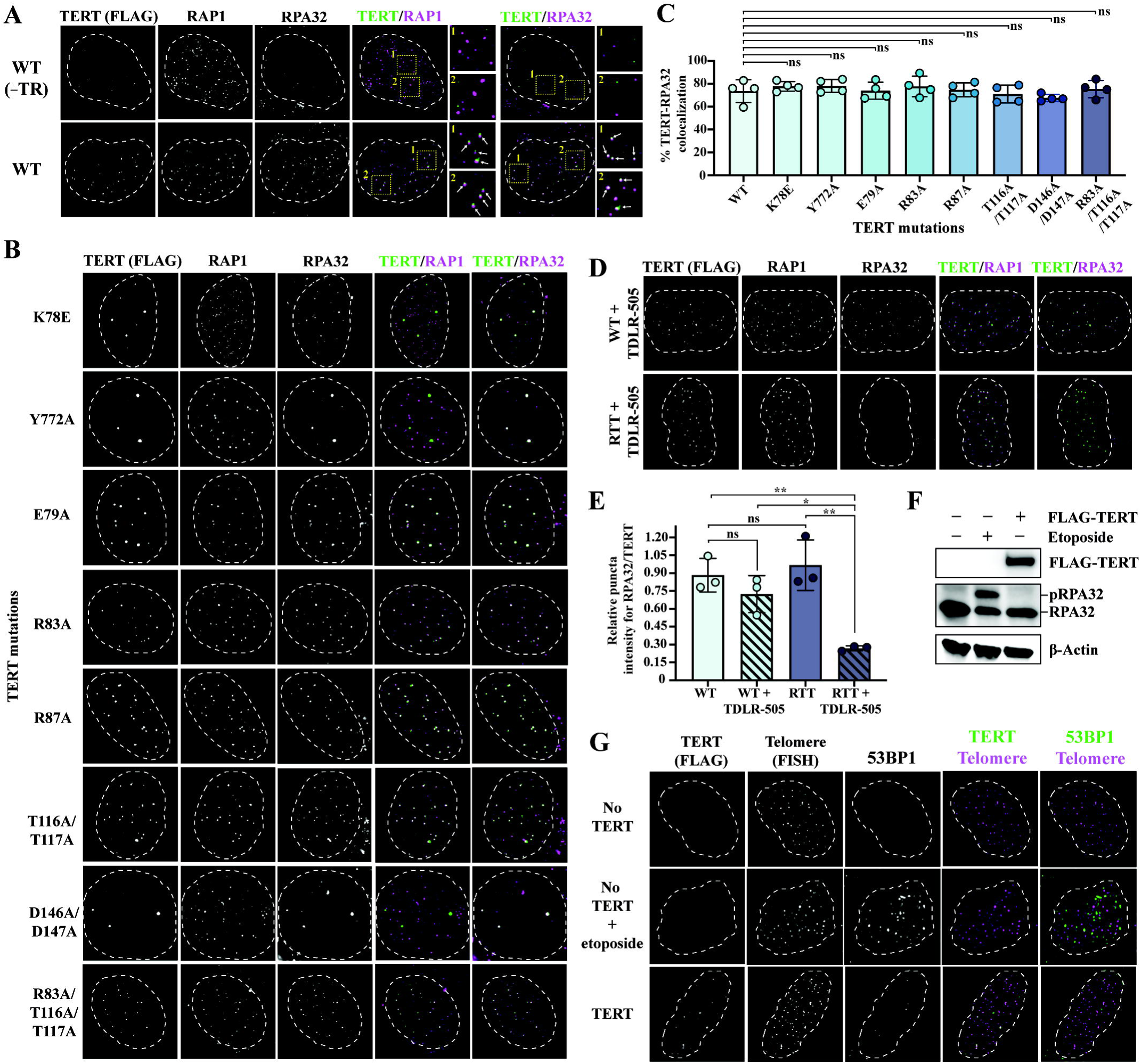
RPA localizes at telomeres via telomerase without triggering a DNA damage response. **(A)** Representative immunofluorescence images of wild-type (WT) TERT with and without telomerase RNA (TR) showing localization of TERT (FLAG), RAP1, and RPA32. Merged images show colocalization of TERT with RAP1 and RPA32. Insets are zoomed in images of representative colocalized puncta, marked by white arrows. **(B)** Immunofluorescence images of TERT mutants (K78E, Y772A, E79A, R83A, R87A, T116A/T117A, D146A/D147A, R83A/T116A/T117A) showing their colocalization with RAP1 and RPA32. **(C)** Quantification of TERT/RPA32 colocalization for WT and TERT mutants. **(D)** Immunofluorescence images comparing WT and RTT (R83A/T116A/T117A) cells treated with RPA ssDNA-binding inhibitor, TDLR-505. **(E)** Quantification of relative RPA32/TERT intensity in WT and RTT cells with and without TDLR-505 treatment. **(F)** Western blot analysis of RPA32 in cells with and without etoposide treatment. FLAG and β-Actin antibodies are used for TERT expression control and loading control respectively. **(G)** Immunofluorescence images showing TERT, telomeres (FISH), and 53BP1 localization in cells with no TERT, TERT with etoposide treatment, and TERT-expressing cells. Nuclei boundaries are demarcated by white dashed lines. Error bars represent standard deviation. ns: not significant (P > 0.05), *: P ≤ 0.05, **: P ≤ 0.01.

To determine if the TERT-RPA32 interaction is crucial for telomerase-dependent RPA localization at telomeres, we performed IF image analysis using various TERT RPA mutants (E79A, R83A, R87A, T116A/T117A, and D146A/D147A). None of these mutations disrupted RPA localization to telomeres (**Fig. 3B & C, and fig. S7A**). To test a more significant disruption, we combined the R83A and T116A/T117A mutations (hereafter referred to as the RTT mutant) but found that even the RTT mutant did not impair RPA localization at telomeres. These results indicate that the TERT-RPA32 interaction, while essential for telomerase RAP stimulation in biochemical assays, appears to be less critical for RPA localization at telomeres, suggesting other factors may also play a role in this process.

Given that these TERT RPA mutations are located away from the known TERT-TPP1 OB domain interaction site (41, 42, 44), we did not expect them to affect telomerase recruitment to telomeres. Our IF imaging experiments confirmed this, except for the D146A/D147A mutant (**Fig. 3B & fig. S7B & C**). The D146A/D147A mutant exhibited a puncta distribution pattern similar to that of the TERT TPT mutants, K78E and Y772A, suggesting that the D146A/D147A mutation impairs telomerase recruitment to telomeres. To validate this, we used IF imaging to investigate whether the mutation caused telomerase to be retained in Cajal bodies, as observed with the TERT TPT mutants (41, 44). Indeed, the mutation led to telomerase retention in Cajal bodies (**fig. S8A**). Interestingly, this mutation also caused endogenous RPA to localize to the Cajal bodies instead of telomeres, suggesting that telomerase associates with RPA at the Cajal bodies as well (**fig. S8A**).

The loss of telomerase recruitment in the TERT D146A/D147A mutant is surprising considering the distance between these mutations and the known TPP1 OB domain interaction sites (41–44) (**fig. S8B & C**). A possible explanation is that in addition to disrupting TERT-RPA32 OB-D domain interactions, these mutations might also perturb TERT-TPP1 OB domain interactions through some structural changes in the TERT TEN domain. Interestingly, these mutations did not significantly impact TPT stimulation of telomerase RAP (**Fig. 2F**), suggesting a complex relationship between TERT’s structural element and its various protein interactions.

To investigate if RPA’s ssDNA-binding ability plays a role in its localization to telomeres, we used the cell-permeable RPA ssDNA-binding inhibitor TDLR-505 (46) in our IF experiments. We found that the inhibitor alone did not affect RPA localization at telomeres in cells expressing WT telomerase (**Fig. 3D & E**). However, when the cells were expressing the TERT RTT mutant, adding the inhibitor resulted in a significant reduction in RPA localization at telomeres as compared to the levels observed in WT or RTT mutant cells (**Fig. 3D & E**). These findings suggest that stable localization of RPA at telomeres depends on RPA binding to both telomerase and the telomeric ssDNA overhang.

Because RPA ssDNA-binding is important for its localization at telomeres, we considered another possibility. We wondered if the lengthening of telomeric overhangs by telomerase overexpression had resulted in more binding sites for RPA, potentially becoming a determinant for RPA localization at telomeres. To test this, we performed IF image analysis in cells expressing a catalytically dead telomerase mutant (TERT D712A) (47). We found that RPA still localized at telomeres at WT levels in cells expressing this mutant (**fig. S9**). This indicates that RPA localization at telomeres is not dependent on binding to longer telomere overhangs that might result from elevated telomerase expression.

Finally, we investigated whether RPA localization at telomeres triggers a DNA damage response (DDR). Previous studies have shown that RPA accumulates at telomeres upon POT1 depletion, activating ATR kinase and leading to RPA32 phosphorylation (48–50). To assess the RPA32 phosphorylation state when telomerase is overexpressed, we performed western blot analysis. Phosphorylation of RPA32 is indicated by a band shift in the western blot (51). Unlike etoposide-treated cells, cells overexpressing telomerase showed no RPA32 phosphorylation (**Fig. 3F**), indicating no ATR activation. To complement our whole cell lysate western blot analysis, we used IF-FISH to examine 53BP1 foci, another DDR marker (49), at telomeres. While etoposide-treated cells displayed numerous foci, telomerase-overexpressing cells showed no 53BP1 foci at telomeres (or the nucleus), similar to the control sample (**Fig. 3G**). This result corroborates our western blot analysis, showing that there was no DDR activation at telomeres.

Collectively, these results demonstrate that telomerase aids RPA localization at telomeres, and this process depends on RPA interactions with both TERT and the telomeric ssDNA overhang. More importantly, our findings indicate that telomerase-mediated RPA localization to telomeres does not activate DDR at telomeres.

## TERT separation-of-function RPA mutants are defective in telomere maintenance in cells

Having established that RPA is localized to telomeres by telomerase, we investigated its physiological function in telomere maintenance. Unlike TPP1-POT1, whose dual roles in recruitment and stimulation complicate functional separation, our SOF TERT RPA mutants (R83A and T116A/T117A) lost their ability to be stimulated by RPA (**Fig. 2F**) but retained their telomerase recruitment function (**Fig. 3B**). This allows us to specifically determine the impact of RPA’s processivity stimulation on telomere maintenance in cultured cells.

We used CRISPR/Cas9 to introduce a WT or mutant TERT expression cassette (K78E, Y772A, E79A, R83A, R87A, T116A/T117A, and D146A/D147A) into the safe harbor genomic locus, *AAVS1*, in HEK293T cells (*52*) (**Fig. 4A**). Polyclonal edited cell lines were verified by genotyping and western blot analysis (**Fig. 4B**). Since endogenous TERT is limited (*53*), exogenous TERT dominates telomerase assembly and leads to a dominant phenotype (44).

**Fig. 4.**
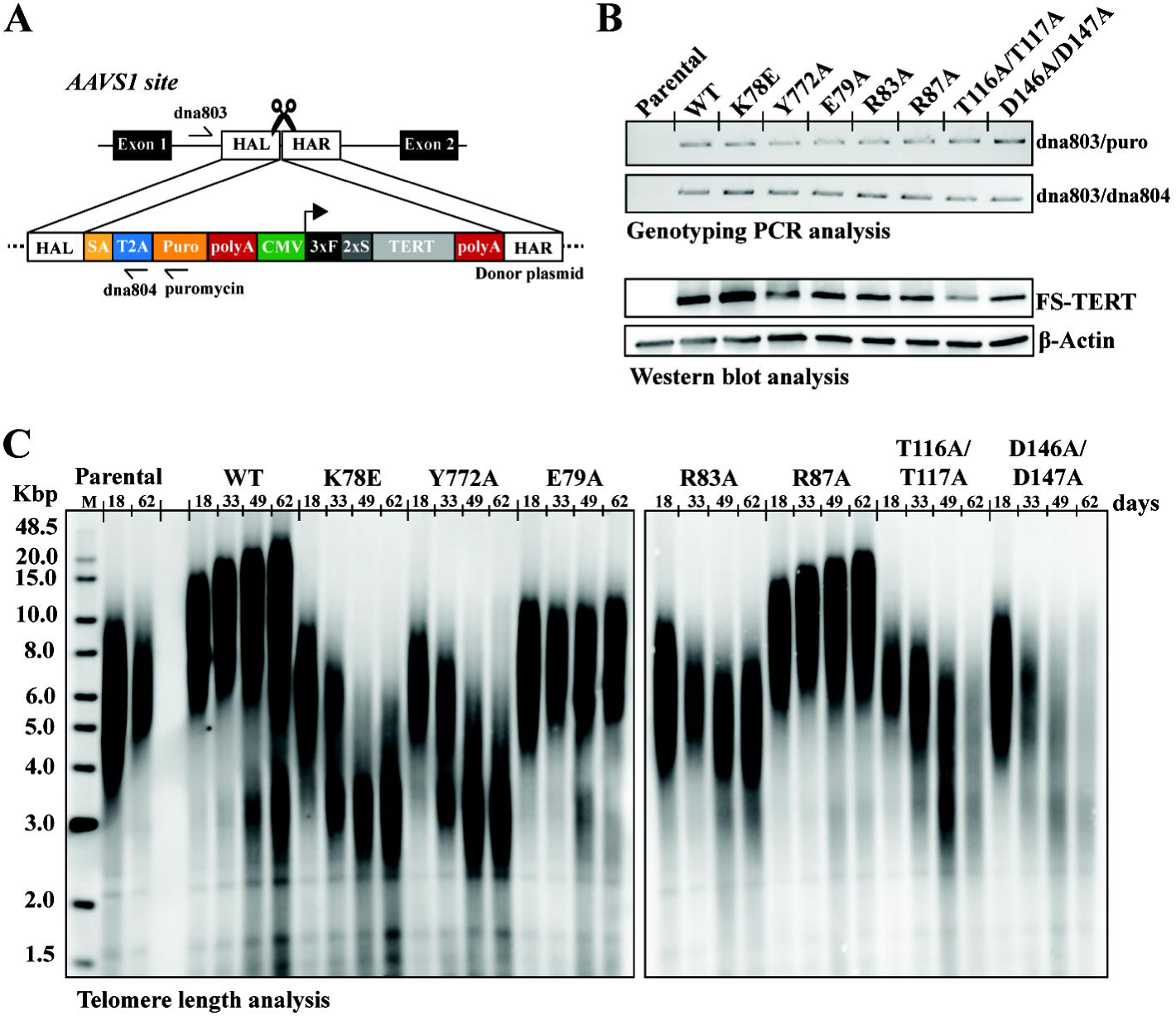
TERT separation-of-function RPA mutants are defective in telomere maintenance in cells. **(A)** Schematic representation of the AAVS1 locus and donor plasmid design for TERT expression cassette genome insertion. The AAVS1 site is located between exons 1 and 2 of the PPP1R12C gene. The donor plasmid contains homology arms (HAL and HAR), Puromycin selection marker, and the TERT cDNA under a CMV promoter control. F: FLAG and S: Strep tag. **(B)** Genotyping PCR analysis confirming successful insertion of TERT constructs at the AAVS1 locus (top two panels) and Western blot analysis showing expression of FLAG-Twin-Strep-tagged TERT (FS-TERT) in selected edited HEK293T cell lines (bottom two panels). β- Actin serves as a loading control. Parental lane serves as FS-tagged TERT expression control. **(C)** Telomere length analysis by Southern blot of parental and edited cell lines (WT, K78E, Y772A, E79A, R83A, R87A, T116A/T117A, D146A/D147A) at different cell culture time points (18, 33, 49, and 62 days). M indicates the molecular weight marker in kilobase pairs (Kbp).

We measured telomere length over time using Southern blot analysis. Exogenous WT TERT expression led to telomere lengthening, as expected due to the elevated telomerase copies (45). The TERT TPT mutant controls (K78E and Y772A) caused telomere attrition (**Fig. 4C**), consistent with their recruitment defect phenotype (**Fig. 3B**) (44, 54). The R83A and T116A/T117A mutants, which retained their telomerase recruitment function (**Fig. 3B**) but cannot support RPA stimulation (**Fig. 2F**), also led to telomere shortening (**Fig. 4C**). This result suggests that RPA telomerase RAP stimulation is essential for telomere maintenance. The D146A/D147A mutant similarly resulted in reduced telomere length (**Fig. 4C**), but this is expected given its telomerase recruitment defect phenotype (**Fig. 3B**). The telomere length phenotypes of the E79A and R87A mutations are consistent with their telomerase direct assay results (**Fig. 2F**), which further underline the correlation between RPA’s telomerase RAP stimulation function and cellular telomere lengthening by telomerase.

Our findings regarding the TERT R83A and T116A/T117A mutations provide important insights into telomere maintenance mechanisms. These mutants retained their ability to be stimulated by TPT but were unable to support telomere maintenance. This suggests that TPP1-POT1-mediated telomerase RAP stimulation may not be the primary factor in telomere lengthening. Instead, our results indicate that RPA plays a crucial role in supporting telomerase extension of telomeres.

## Connections to short telomeres human diseases

We next investigated whether our findings on RPA-TERT interactions could provide insights into telomere-related diseases. We noted that several short telomere disease mutations, including TERT R83P (55), F115L (56), and T116I (57), occur within the predicted interaction site between TERT and the RPA32 OB-D domain (**Fig. 5A)**. To assess the functional impact of these mutations, we performed direct telomerase assays comparing wild-type TERT with these disease-associated mutants.

**Fig. 5.**
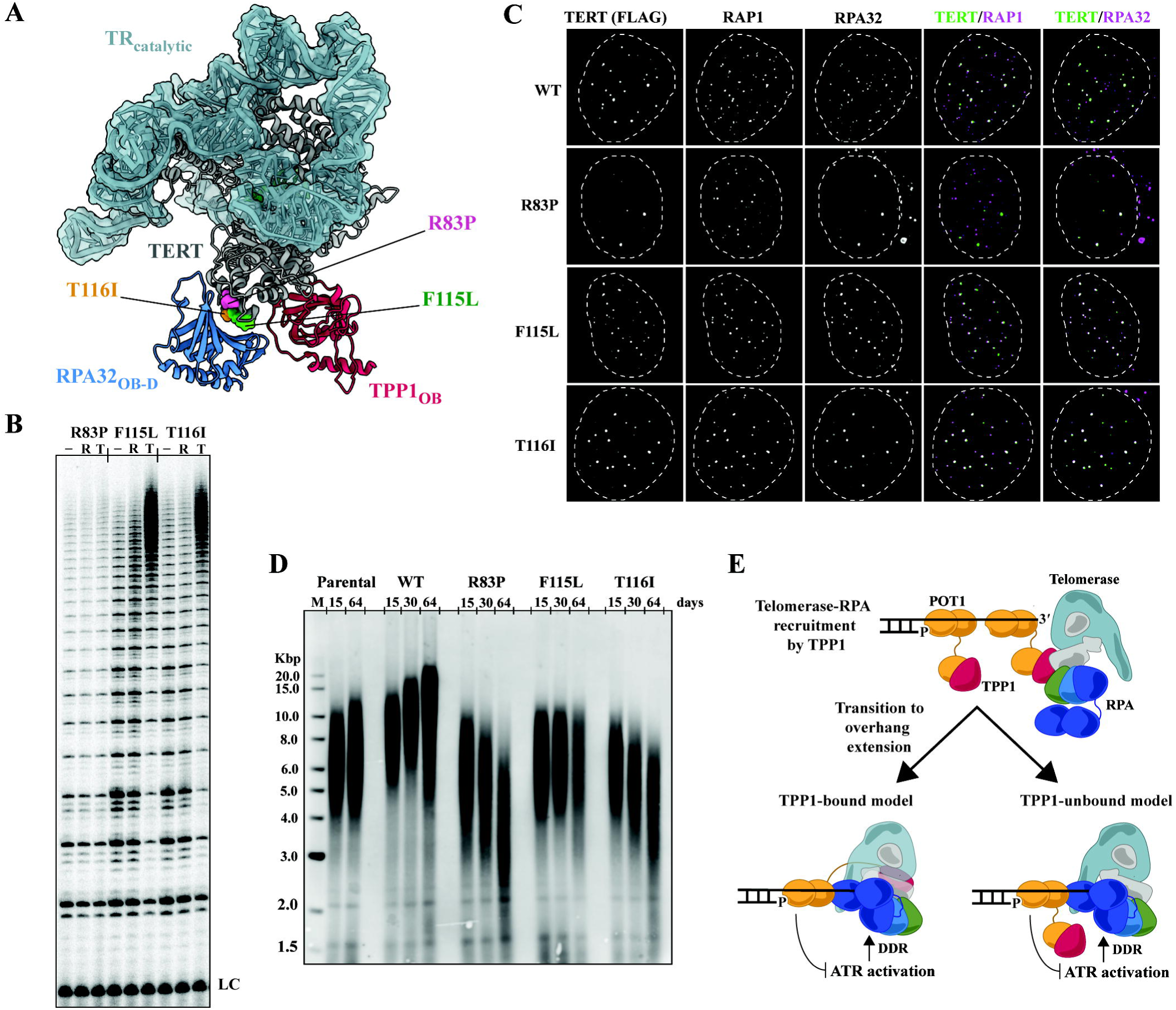
Loss in telomerase RAP stimulation by RPA provides an explanation for short telomere disease TERT mutations. **(A)** Short telomere disease mutations mapped onto the AlphaFold2 multimer model of the telomerase-RPA32_OB-D_-TPP1_OB_ complex. The disease mutations R83P, F115L, and T116I are highlighted in pink, green, and orange, respectively. **(B)** Direct telomerase assays comparing wild-type (WT) TERT with the disease mutations (R83P, F115L, T116I) in the presence or absence of RPA (R) and TPT (T). LC denotes loading control. **(C)** Immunofluorescence images of WT TERT and the mutants showing localization of TERT (FLAG), RAP1, and RPA32. Merged images show colocalization of TERT with RAP1 and RPA32. Nuclei boundaries are demarcated by white dashed lines in panel **C**. **(D)** Telomere length analysis by Southern blot of parental cells and edited cell lines (WT, R83P, F115L, T116I) at different time points (15, 30, and 64 days). M indicates the molecular weight marker in kilobase pairs (Kbp). **(E)** Proposed updated model for telomerase regulation by TPP1-POT1 and RPA. The telomerase-RPA complex is recruitment by TPP1 and then transits to telomerase overhang extension mediated by RPA. Two possible models for the latter step are illustrated; TPP1-bound and TPP1-unbound. In both cases, RPA is not phosphorylated by the ATR kinase, as the activation of the kinase is suppressed by POT1 binding to the recessed telomere 5’ end. Thus, DNA damage response is not triggered at telomeres in the presence of RPA. The cartoon models are not drawn to scale and the overhang is illustrated short for simplicity.

Our results show that these mutations significantly impair RPA-mediated RAP stimulation (**Fig. 5B**). Interestingly, we observe distinct effects among the mutants. The R83P mutation affects both RPA and TPT stimulation, likely due to destabilization of the α4 helix (residues 76-92) (58) in the TEN domain that houses K78, a residue critical for TPP1 interaction (44). In contrast, F115L and T116I mutations selectively disrupt RPA stimulation without affecting TPT-mediated stimulation (**Fig. 5B**). These results identify F115L and T116I as separation-of-function mutants specifically affecting RPA-mediated telomerase stimulation.

To further characterize these disease-associated mutations, we examined their effects in cultured cells. IF imaging revealed that the TERT R83P mutant failed to localize to telomeres (**Fig. 5C**), consistent with its inability to be stimulated by TPT in vitro. In contrast, the F115L and T116I mutations did not affect telomerase recruitment, mirroring the behavior of our previously identified separation-of-function TERT RPA mutants, R83A and T116A/T117A (**Fig. 3B**). Analysis of telomere length maintenance in HEK293T cells expressing these mutants revealed phenotypes consistent with their clinical manifestations (**Fig. 5D).** While WT TERT expression led to telomere lengthening as previously observed, all three disease-associated mutations resulted in either no telomere growth or shortening. The TERT R83P mutant showed the most severe phenotype with progressive telomere shortening, aligning with its recruitment defect. The T116I mutant, despite localizing to telomeres, also led to telomere shortening, demonstrating that loss of RPA-mediated stimulation significantly impairs telomerase activity in vivo. Interestingly, the F115L mutant maintained telomere length but did not induce lengthening like wild-type TERT, indicating a partial loss of telomerase activity.

These results demonstrate a spectrum of telomere maintenance defects among the disease-associated mutations, with severity correlating to their biochemical defects. The phenotypes of the TERT F115L and T116I mutations closely resemble those of our TERT RPA separation-of-function mutants, suggesting that disruption of TERT-RPA interaction contributes to the pathogenesis of associated short telomere diseases. Collectively, these findings underscore the physiological importance of TERT-RPA interaction for telomere maintenance and provide a potential mechanistic explanation for how these disease-associated mutations lead to telomere dysfunction.

## Discussion

Our discovery that human Replication Protein A (RPA) is a crucial telomerase processivity factor reshapes our understanding of telomere maintenance. We propose an updated model (**Fig. 5F**) integrating RPA’s role with established mechanisms. In this model, telomerase is recruited to telomeres via TPP1, followed by RPA association. POT1 protects the ds-ssDNA telomeric junction (59), suppressing DNA damage response activation of RPA. The telomerase-RPA complex then engages the telomeric ssDNA 3’ overhang, promoting processive telomere repeat addition.

Interestingly, this role of RPA was presaged by the structure of the Tetrahymena telomerase holoenzyme, where the N-terminal domain of the Teb2 subunit of the TEB complex, an RPA homolog in ciliates, binds to a similar TERT position (21) as the AlphaFold2-predicted RPA32 OB-D-TERT interaction site. An intact TEB complex is required for optimal telomerase activity (21), similar to our findings here (**Fig. 1**). This evolutionary conservation underscores the fundamental importance of RPA-like proteins in telomerase function across diverse eukaryotes.

Our AlphaFold2 model and the human telomerase-TPP1-POT1 co-complex cryo-EM structure (42) indicate that POT1 could clash with the RPA32 OB-D domain when both are bound to telomerase (**fig. S10**). This raises the question of how these conflicting interactions can be reconciled. Based on our model (**Fig. 5F),** POT1 needs to disengage from telomerase for RPA to take over to facilitate telomerase addition of telomeric DNA repeats. Thus, we speculate that the telomerase-TPP1-POT1 cryo-EM structure may represent a state before the handover. This handover process may be mediated by TERRA and hnRNPA1 (50, 60).

Our work has implications for understanding both telomere-related diseases and neo-telomere formation. The identification of disease-associated TERT mutations that specifically disrupt RPA-mediated stimulation provides a new mechanistic link to short-telomere syndromes. This discovery offers new avenues for potential therapeutic interventions targeting the RPA- telomerase interaction. Additionally, the ability of RPA to promote telomerase processivity without triggering a DNA damage response suggests a potential role in aiding telomerase to add internal telomeric repeats at resected double-strand breaks (61, 62).

The use of AlphaFold2 modeling proved invaluable in guiding our structure-function analyses, further demonstrating its power in studying complex macromolecular assemblies (63), as has been shown by other recent studies (64). Future research should focus on elucidating the molecular details of telomerase-shelterin-RPA interplay during the transition from telomerase recruitment to telomere overhang extension, and investigating the dynamics between POT1, RPA, and telomerase at telomeres.

## Acknowledgments

We thank Dr. Vicki Lundblad, Dr. Deborah Wuttke, Dr. Michael Stone, Dr. Jens Schmidt, Dr. Scott Cohen, and Dr. Tracy Bryan for their constructive suggestions and discussions. We also appreciate the valuable input from Dr. Tom Record and colleagues in the Lim and Huang labs. We also thank Dr. Kelly Nguyen for her generosity in sharing her hTR expression plasmid. We acknowledge the COSMIC^2^ platform for supporting our initial AlphaFold computational work. Finally, we extend our gratitude to Mr. Jeremy Herr and the departmental IT team for their assistance in setting up our computational workflow.

## Funding

Support for this research was provided to C.L. by the National Institutes of Health, the National Institute of General Medical Sciences (R01GM153806 and DP2GM150023) and the University of Wisconsin–Madison, Office of the Vice-Chancellor for Research and Graduate Education with funding from the Wisconsin Alumni Research Foundation and the Department of Biochemistry. X.H. acknowledges the support from the Hirschfelder Professorship Fund. In addition, M.S.O and K.M.A are supported by a NIH T32 predoctoral fellowship (T32GM130550).

## Author contributions

Conceptualization: CL. Methodology: SA, XL, VS, MSO, BLC, VRT, XH, and CL. Investigation: SA, XL, VS, MSO, BLC, VRT, QH, KMA, and CL. Visualization: VS and CL. Funding acquisition: CL and XH. Project administration: CL. Supervision: SA, CL and XH. Writing – original draft: CL and SA. Writing – review & editing: CL, SA, XL, VS, MSO, BLC, VRT, QH, KMA, and XH.

## Competing interests

The authors declare no competing interests.

## Data and materials availability

Plasmids and cell lines generated are available upon request. All data are available in the main text or supplementary materials.

## Supplementary Materials

Materials and Methods

Figs. S1 to S10

Tables S1

References (65-84)

## Notes

### Competing Interest Statement

The authors have declared no competing interest.

## References

1. R. J. O’Sullivan, J. Karlseder, Telomeres: Protecting chromosomes against genome instability. [Preprint] (2010). 10.1038/nrm2848.

2. C. J. Lim, T. R. Cech, Shaping human telomeres: from shelterin and CST complexes to telomeric chromatin organization. Nat Rev Mol Cell Biol 22, 283–298 (2021).

3. H. Takai, V. Aria, P. Borges, J. T. P. Yeeles, T. De Lange, CST–polymerase α-primase solves a second telomere end-replication problem. Nature, doi: 10.1038/s41586-024-07137-1 (2024).

4. R. K. Moyzis, J. M. Buckingham, L. S. Cram, M. Dani, L. L. Deaven, M. D. Jones, J. Meyne, R. L. Ratliff, J. R. Wu, A highly conserved repetitive DNA sequence, (TTAGGG)(n), present at the telomeres of human chromosomes. Proc Natl Acad Sci U S A 85 (1988).

5. C. W. Greider, E. H. Blackburn, The telomere terminal transferase of tetrahymena is a ribonucleoprotein enzyme with two kinds of primer specificity. Cell 51 (1987).

6. C. W. Greider, E. H. Blackburn, A telomeric sequence in the RNA of Tetrahymena telomerase required for telomere repeat synthesis. Nature 337 (1989).

7. J. Lingner, T. R. Hughes, A. Shevchenko, M. Mann, V. Lundblad, T. R. Cech, Reverse transcriptase motifs in the catalytic subunit of telomerase. Science (1979) 276 (1997).

8. T. M. Nakamura, G. B. Morin, K. B. Chapman, S. L. Weinrich, W. H. Andrews, J. Lingner, C. B. Harley, T. R. Cech, Telomerase catalytic subunit homologs from fission yeast and human. Science (1979) 277 (1997).

9. Q. He, C. J. Lim, Models for human telomere C-strand fill-in by CST–Polα-primase. [Preprint] (2023). 10.1016/j.tibs.2023.07.008.

10. S. W. Cai, T. de Lange, CST–Polα/Primase: the second telomere maintenance machine. [Preprint] (2023). 10.1101/GAD.350479.123.

11. H. Xin, D. Liu, M. Wan, A. Safari, H. Kim, W. Sun, M. S. O’Connor, Z. Songyang, TPP1 is a homologue of ciliate TEBP-β and interacts with POT1 to recruit telomerase. Nature 445 (2007).

12. E. Abreu, E. Aritonovska, P. Reichenbach, G. Cristofari, B. Culp, R. M. Terns, J. Lingner, M. P. Terns, TIN2-Tethered TPP1 Recruits Human Telomerase to Telomeres In Vivo. Mol Cell Biol 30 (2010).

13. A. M. Tejera, M. Stagno d’Alcontres, M. Thanasoula, R. M. Marion, P. Martinez, C. Liao, J. M. Flores, M. Tarsounas, M. A. Blasco, TPP1 is required for TERT recruitment, telomere elongation during nuclear reprogramming, and normal skin development in mice. Dev Cell 18 (2010).

14. F. L. Zhong, L. F. Z. Batista, A. Freund, M. F. Pech, A. S. Venteicher, S. E. Artandi, TPP1 OB-fold domain controls telomere maintenance by recruiting telomerase to chromosome ends. Cell 150 (2012).

15. J. Nandakumar, C. F. Bell, I. Weidenfeld, A. J. Zaug, L. A. Leinwand, T. R. Cech, The TEL patch of telomere protein TPP1 mediates telomerase recruitment and processivity. Nature 492 (2012).

16. A. N. Sexton, D. T. Youmans, K. Collins, Specificity requirements for human telomere protein interaction with telomerase holoenzyme. Journal of Biological Chemistry 287 (2012).

17. F. Wang, E. R. Podell, A. J. Zaug, Y. Yang, P. Baciu, T. R. Cech, M. Lei, The POT1-TPP1 telomere complex is a telomerase processivity factor. Nature 445 (2007).

18. C. M. Latrick, T. R. Cech, POT1-TPP1 enhances telomerase processivity by slowing primer dissociation and aiding translocation. EMBO Journal 29 (2010).

19. M. S. Wold, Replication protein A: A heterotrimeric, single-stranded DNA-binding protein required for eukaryotic DNA metabolism. [Preprint] (1997). 10.1146/annurev.biochem.66.1.61.

20. R. Dueva, G. Iliakis, Replication protein A: a multifunctional protein with roles in DNA replication, repair and beyond. [Preprint] (2020). 10.1093/narcan/zcaa022.

21. J. Jiang, H. Chan, D. D. Cash, E. J. Miracco, R. R. O. Loo, H. E. Upton, D. Cascio, R. O. B. Johnson, K. Collins, J. A. Loo, Z. H. Zhou, J. Feigon, Structure of Tetrahymena telomerase reveals previously unknown subunits, functions, and interactions. Science (1979) 350 (2015).

22. H. E. Upton, H. Chan, J. Feigon, K. Collins, Shared subunits of tetrahymena telomerase holoenzyme and replication protein a have different functions in different cellular complexes. Journal of Biological Chemistry 292 (2017).

23. Z. Zeng, B. Min, J. Huang, K. Hong, Y. Yang, K. Collins, M. Lei, Structural basis for Tetrahymena telomerase processivity factor Teb1 binding to single-stranded telomeric-repeat DNA. Proc Natl Acad Sci U S A 108 (2011).

24. H. E. Upton, K. Hong, K. Collins, Direct Single-Stranded DNA Binding by Teb1 Mediates the Recruitment of Tetrahymena thermophila Telomerase to Telomeres. Mol Cell Biol 34 (2014).

25. P. Luciano, S. Coulon, V. Faure, Y. Corda, J. Bos, S. J. Brill, E. Gilson, M. N. Simon, V. Géli, RPA facilitates telomerase activity at chromosome ends in budding and fission yeasts. EMBO Journal 31 (2012).

26. V. Schramke, P. Luciano, V. Brevet, S. Guillot, Y. Corda, M. P. Longhese, E. Gilson, V. Géli, RPA regulates telomerase action by providing Est1p access to chromosome ends. Nat Genet 36 (2004).

27. H. Chan, Y. Wang, J. Feigon, Progress in Human and Tetrahymena Telomerase Structure Determination. [Preprint] (2017). 10.1146/annurev-biophys-062215-011140.

28. Y. V. Surovtseva, D. Churikov, K. A. Boltz, X. Song, J. C. Lamb, R. Warrington, K. Leehy, M. Heacock, C. M. Price, D. E. Shippen, Conserved Telomere Maintenance Component 1 Interacts with STN1 and Maintains Chromosome Ends in Higher Eukaryotes. Mol Cell 36 (2009).

29. Y. Miyake, M. Nakamura, A. Nabetani, S. Shimamura, M. Tamura, S. Yonehara, M. Saito, F. Ishikawa, RPA-like Mammalian Ctc1-Stn1-Ten1 Complex Binds to Single-Stranded DNA and Protects Telomeres Independently of the Pot1 Pathway. Mol Cell 36 (2009).

30. C. J. Lim, A. T. Barbour, A. J. Zaug, K. J. Goodrich, A. E. McKay, D. S. Wuttke, T. R. Cech, The structure of human CST reveals a decameric assembly bound to telomeric DNA. Science (1979) 368, 1081–1085 (2020).

31. L. Y. Chen, S. Redon, J. Lingner, The human CST complex is a terminator of telomerase activity. Nature 488 (2012).

32. A. J. Zaug, C. J. Lim, C. L. Olson, M. T. Carilli, K. J. Goodrich, D. S. Wuttke, T. R. Cech, CST does not evict elongating telomerase but prevents initiation by ssDNA binding. Nucleic Acids Res 49, 11653–11665 (2021).

33. E. Bochkareva, S. Korolev, S. P. Lees-Miller, A. Bochkarev, Structure of the RPA trimerization core and its role in the multistep DNA-binding mechanism of RPA. EMBO Journal 21 (2002).

34. S. K. Binz, M. S. Wold, Regulatory functions of the N-terminal domain of the 70-kDa subunit of replication protein A (RPA). Journal of Biological Chemistry 283 (2008).

35. X. V. Gomes, M. S. Wold, Functional domains of the 70-kilodalton subunit of human replication protein A. Biochemistry 35 (1996).

36. A. Bochkarev, R. A. Pfuetzner, A. M. Edwards, L. Frappier, Structure of the single-stranded-DNA-binding domain of replication protein A bound to DNA. Nature 385 (1997).

37. C. Kim, M. S. Wold, Recombinant Human Replication Protein A Binds to Polynucleotides with Low Cooperativity. Biochemistry 34 (1995).

38. P. Baumann, T. R. Cech, Pot1, the putative telomere end-binding protein in fission yeast and humans. Science (1979) 292 (2001).

39. R. A. Wu, K. Collins, Sequence specificity of human telomerase. [Preprint] (2014). 10.1073/pnas.1411276111.

40. J. Jumper, R. Evans, A. Pritzel, T. Green, M. Figurnov, O. Ronneberger, K. Tunyasuvunakool, R. Bates, A. Žídek, A. Potapenko, A. Bridgland, C. Meyer, S. A. A. Kohl, A. J. Ballard, A. Cowie, B. Romera-Paredes, S. Nikolov, R. Jain, J. Adler, T. Back, S. Petersen, D. Reiman, E. Clancy, M. Zielinski, M. Steinegger, M. Pacholska, T. Berghammer, S. Bodenstein, D. Silver, O. Vinyals, A. W. Senior, K. Kavukcuoglu, P. Kohli, D. Hassabis, Highly accurate protein structure prediction with AlphaFold. Nature 596 (2021).

41. B. Liu, Y. He, Y. Wang, H. Song, Z. H. Zhou, J. Feigon, Structure of active human telomerase with telomere shelterin protein TPP1. Nature 604 (2022).

42. Z. Sekne, G. E. Ghanim, A.-M. M. Van Roon, T. H. D. Nguyen, Structural basis of human telomerase recruitment by TPP1-POT1. Science (1979) 375, 1173–1176 (2022).

43. S. Padmanaban, V. M. Tesmer, J. Nandakumar, Interaction hub critical for telomerase recruitment and primer-template handling for catalysis. Life Sci Alliance 6 (2023).

44. J. C. Schmidt, A. B. Dalby, T. R. Cech, Identification of human TERT elements necessary for telomerase recruitment to telomeres. Elife 3 (2014).

45. T. A. Wieser, D. S. Wuttke, Replication Protein A Utilizes Differential Engagement of Its DNA-Binding Domains to Bind Biologically Relevant ssDNAs in Diverse Binding Modes. Biochemistry 61 (2022).

46. A. K. Mishra, S. S. Dormi, A. M. Turchi, D. S. Woods, J. J. Turchi, Chemical inhibitor targeting the replication protein A-DNA interaction increases the efficacy of Pt-based chemotherapy in lung and ovarian cancer. Biochem Pharmacol 93 (2015).

47. T. L. Beattie, W. Zhou, M. O. Robinson, L. Harrington, Functional Multimerization of the Human Telomerase Reverse Transcriptase. Mol Cell Biol 21 (2001).

48. L. Zou, S. J. Elledge, Sensing DNA damage through ATRIP recognition of RPA-ssDNA complexes. Science (1979) 300 (2003).

49. E. L. Denchi, T. De Lange, Protection of telomeres through independent control of ATM and ATR by TRF2 and POT1. Nature 448 (2007).

50. R. L. Flynn, R. C. Centore, R. J. O’Sullivan, R. Rai, A. Tse, Z. Songyang, S. Chang, J. Karlseder, L. Zou, TERRA and hnRNPA1 orchestrate an RPA-to-POT1 switch on telomeric single-stranded DNA. Nature 471 (2011).

51. Salah-ud-Din, S. J. Brill, M. P. Fairman, B. Stillman, Cell-cycle-regulated phosphorylation of DNA replication factor A from human and yeast cells. Genes Dev 4 (1990).

52. R. C. DeKelver, V. M. Choi, E. A. Moehle, D. E. Paschon, D. Hockemeyer, S. H. Meijsing, Y. Sancak, X. Cui, E. J. Steine, J. C. Miller, P. Tam, V. V. Bartsevich, X. Meng, I. Rupniewski, S. M. Gopalan, H. C. Sun, K. J. Pitz, J. M. Rock, L. Zhang, G. D. Davis, E. J. Rebar, I. M. Cheeseman, K. R. Yamamoto, D. M. Sabatini, R. Jaenisch, P. D. Gregory, F. D. Urnov, Functional genomics, proteomics, and regulatory DNA analysis in isogenic settings using zinc finger nuclease-driven transgenesis into a safe harbor locus in the human genome. Genome Res 20 (2010).

53. L. Xi, T. R. Cech, Inventory of telomerase components in human cells reveals multiple subpopulations of hTR and hTERT. Nucleic Acids Res 42 (2014).

54. J. C. Schmidt, A. J. Zaug, T. R. Cech, Live Cell Imaging Reveals the Dynamics of Telomerase Recruitment to Telomeres. Cell 166 (2016).

55. T. J. Vulliamy, M. J. Kirwan, R. Beswick, U. Hossain, C. Baqai, A. Ratcliffe, J. Marsh, A. Walne, I. Dokal, Differences in disease severity but similar telomere lengths in genetic subgroups of patients with telomerase and shelterin mutations. [Preprint] (2011). 10.1371/journal.pone.0024383.

56. K. E. Schratz, L. Haley, S. K. Danoff, A. L. Blackford, A. E. DeZern, C. D. Gocke, A. S. Duffield, M. Armanios, Cancer spectrum and outcomes in the Mendelian short telomere syndromes. Blood 135 (2020).

57. J. D. Podlevsky, C. J. Bley, R. V. Omana, X. Qi, J. L. Chen, The Telomerase Database. Nucleic Acids Res 36 (2008).

58. Y. Wang, M. Gallagher-Jones, L. Sušac, H. Song, J. Feigon, A structurally conserved human and Tetrahymena telomerase catalytic core. Proc Natl Acad Sci U S A 117 (2020).

59. V. M. Tesmer, K. A. Brenner, J. Nandakumar, Human POT1 protects the telomeric ds-ss DNA junction by capping the 5′ end of the chromosome. Science (1979) 381 (2023).

60. S. L. Granger, R. Sharma, V. Kaushik, M. Razzaghi, M. Honda, D. S. Bhat, M. Wlodarski, E. Antony, M. Spies, Human hnRNPA1 reorganizes telomere-bound Replication Protein A. bioRxiv (2023).

61. A. O. M. Wilkie, J. Lamb, P. C. Harris, R. D. Finney, D. R. Higgs, A truncated human chromosome 16 associated with α thalassaemia is stabilized by addition of telomeric repeat (TTAGGG)n. Nature 346 (1990).

62. C. G. Kinzig, G. Zakusilo, K. K. Takai, L. R. Myler, T. de Lange, ATR blocks telomerase from converting DNA breaks into telomeres. Science (1979) 383 (2024).

63. Y. Lim, L. Tamayo-Orrego, E. Schmid, Z. Tarnauskaite, O. V. Kochenova, R. Gruar, S. Muramatsu, L. Lynch, A. V. Schlie, P. L. Carroll, G. Chistol, M. A. M. Reijns, M. T. Kanemaki, A. P. Jackson, J. C. Walter, In silico protein interaction screening uncovers DONSON’s role in replication initiation. Science (1979) 381 (2023).

64. E. A. Campbell, H. Walden, J. C. Walter, A. K. Shukla, M. Beck, L. A. Passmore, H. E. Xu, AlphaFold: Research accelerator and hypothesis generator. [Preprint] (2024). 10.1016/j.molcel.2023.12.035.

