## Supplementary Information for "Human Replication Protein A complex is a Telomerase Processivity Factor Essential for Telomere Maintenance"

**This PDF file includes:**

Materials and Methods
Figs. S1 to S10
Table S1
References (65-84)

### Materials and Methods

#### Molecular cloning

All the cloning is performed in chemically competent *E. coli* XL1-Blue cells (New England Biolabs). The constructs were either generated by site-directed mutagenesis or Hi-fi Assembly reactions (New England Biolabs). The primers used for subsequent construct generation are listed in **Supplementary Table S1**. The assembled constructs were sequence verified using Oxford Nanopore Sequencing (Plasmidsaurus).

#### Expression and purification of human telomerase

HEK293T cells (ATCC) were transfected with a modified version of pVan107-hTERT (1) having a 3xFLAG tag followed by a Twin-Strep Tag at the N-terminal of TERT and pCDNA-U3-TR-HDV (a kind gift from Dr. Kelly Nguyen) in a 1:3 mass ratio using JetPrime reagent (Polyplus). The cells were collected after 48 h of transfection and lysed by resuspending in CHAPS lysis buffer (10 mM HEPES pH 7.5, 1 mM $MgCl_2$, 1 mM EGTA, 0.5% (v/w) CHAPS, 10% Glycerol, 5 mM TCEP, 1 mM PMSF). The lysate was centrifuged at 17,000 x g for 20 min and the supernatant was added to anti-DYKDDDDK G1 Affinity Resin (Genscript) and incubated at 4 °C for 2 h under constant rotation. The beads were washed with 15 CV of Telomerase Buffer (50 mM Tris-HCl pH 8.0, 50 mM KCl, 1 mM MgCl2, 1 mM TCEP, 30% Glycerol). The protein was eluted with 2 CV of telomerase Elution Buffer [50 mM Tris-HCl pH 8.0, 50 mM KCl, 1 mM $MgCl_2$, 1 mM TCEP, 30% Glycerol, 0.5 mg $mL^{-1}$ 3xFLAG peptide (APExBIO)]. The elute was concentrated using Amicon Ultra 30 KDa MWCO centrifugal tubes (Millipore) and snap-frozen in small aliquots using liquid $N_2$ and stored in –80 °C.

The presence of TERT in purified telomerase was confirmed and quantified using western blot. The purified telomerase was run on SDS-PAGE gel against a 3xFLAG tagged protein standard and then transferred to nitrocellulose membrane through a semi-dry transfer module. The membrane was blocked by StartingBlock$^{TM}$ Blocking Buffer (ThermoFisher Scientific) for 1 h at room temperature. This was followed by overnight incubation with monoclonal anti-FLAG M2 antibody (Sigma-Aldrich) at recommended concentration overnight at 4 °C. The membrane was washed three times in 1x PBST (with 0.5% Tween-20) and was visualized using SuperSignal West Pico PLUS chemiluminescence substrate (ThermoFisher Scientific).

The presence of TR (Telomerase RNA) in purified telomerase was confirmed and quantified using Dot blot assay. Purified telomerase aliquots were added with Proteinase K and RNaseOUT (Invitrogen) and incubated at 37 °C for 1 h. It was added to an equal amount of Formamide Loading dye and heated at 95 °C for 10 min. This was then dotted to Hybond N+ membrane (Cytiva) along with in vitro transcribed hTR as a standard and crosslinked using UV crosslinker. The membrane was hybridized for 1 h at 42 °C using ULTRAhyb Ultrasensitive Hybridization Buffer (Invitrogen) and then incubated overnight at 42 °C with radioactive P32-labelled hTR DNA probes (mentioned in **Supplementary Table S1**). The membrane was washed with Buffer W1 (2x SSC, 0.1% SDS) and Buffer W2 (0.1x SSC, 0.1% SDS) twice each for 15 minutes at 42 °C. The membrane was exposed to a storage phosphor screen. After adequate exposure, the gels were imaged on a Typhoon FLA9000 gel imager (Cytiva). The hTR quantification was used as the final concentration of purified telomerase.

**Expression and purification of recombinant human RPA proteins and complexes**

RPA and its single amino acid mutants (T88A, W107A, H131A) were purified using the protocol from Wold lab (2). The plasmid p11d-tRPA (Addgene, catalog #102613), and its respective mutant plasmids were transformed into *E. coli* BL21(DE3) cells (New England Biolabs). Single colony was picked and left to grow in 1 L of LB media overnight at 37 °C without shaking. They were grown the next day and induced at O.D. (600) of 0.6 by adding 0.3 mM IPTG. The cells were grown overnight at 18 °C and collected the next day. The cells were resuspended in Buffer A (20 mM HEPES, 10% Glycerol, 0.01% NP-40, 25 μM EDTA, 1 mM DTT) and sonicated for lysis. The lysate was centrifuged at 35,000 x g for 45 min. The supernatant was loaded into the HiTrap-Blue HP column (Cytiva) and subsequently washed with Buffer A containing 50 mM KCl, 800 mM KCl and 400 mM NaSCN for 5 Column Volume (CV) each. The RPA was eluted with Buffer A containing 1.5 M NaSCN. The elute fraction peaks were collected and loaded into the HiTrap Desalting column (Cytiva) to get rid of excess salts and elute protein fraction peaks were subsequently loaded into HiTrap Q column. The Q column was washed with wash buffers having salt concentration 50 mM KCl, 80 mM KCl and 200 mM KCl. The protein was eluted by a gradient of 200-600 mM KCl. The protein peaks were analyzed in SDS-PAGE gel and the pure fractions were pooled together and concentrated using Amicon Ultra 10 KDa MWCO centrifugal tubes (Millipore) to get RPA protein.

For the OB-domain deletion mutants of RPA, a 3xDYKDDDDK tag was added to RPA70 and a 6xHIS tag was added to RPA14. The complexes were purified using tandem affinity purification. The respective mutant plasmids were transformed and induced similarly as mentioned in the Wold Lab protocol (2). The cells were collected and resuspended in Buffer A and sonicated for lysis. The lysate was centrifuged at 35,000 x g for 45 min and the supernatant was collected. The salt concentration in lysate was now increased to make a final concentration of 300 mM NaCl. The lysate was now incubated with Ni-NTA Agarose resin (Qiagen) for 2 h. The beads were washed with 30 CV Wash buffer (20 mM HEPES, 300 mM NaCl, 15 mM Imidazole, 0.01% NP-40, 25 μM EDTA, 1 mM DTT) and eluted with 10 CV Elution Buffer having 300 mM Imidazole. The elute was incubated with anti-DYKDDDDK G1 Affinity Resin (Genscript) overnight. The beads were washed with Wash Buffer (20 mM HEPES, 150 mM NaCl, 0.01% NP-40, 25 μM EDTA, 1 mM DTT) and eluted with Elution Buffer [20 mM HEPES, 150 mM NaCl, 0.01% NP-40, 25 μM EDTA, 1 mM DTT, 0.5 mg $mL^{-1}$ 3x FLAG peptides (APExBIO)]

RPA14-32 complex and RPA70 were purified from *T. ni* insect cells (Expression Systems). The genes with a 3xFLAG tag on RPA70 and 6xHIS tag on both RPA 14 and RPA32 were cloned into two separate pBac4x vector (one having RPA70 and other having RPA14-32). The baculoviruses expressing these proteins were made using the flashBAC ULTRA system (Mirus Bio). All viruses were made using *S. frugiperda* (SF9) cells (Invitrogen). For RPA 14-32, both the viruses encoding RPA14-32 and RPA70 were used to infect one or two liters of *T. ni* cells (Expression Systems). The cells were collected 68 hours after infection and resuspended in lysis buffer [20 mM HEPES, 300 mM NaCl, 15 mM Imidazole, 1 mM DTT, 1x Protease Inhibitor (GenDEPOT)] and then sonicated for lysis. The lysate was centrifuged (35,000 x g, 45 min) and supernatant was collected. Tandem affinity purification was done like OB-domain mutants mentioned above using Ni-NTA and Anti-FLAG tag resin using same buffers mentioned above to get some full-length RPA. The Flowthrough from the anti-Flag beads containing excess RPA14-32 was loaded into Superdex 75 column (Cytiva) with a constant flow of SEC Buffer (20 mM HEPES, 150 mM NaCl, 1 mM DTT). The protein peak fractions were collected and analyzed using SDS-PAGE gel. Fractions with suitable purity were pooled together and concentrated to get RPA14-32. For RPA70, the Baculovirus expressing RPA70 was used to infect one or two liters of *T. ni* cells (Expression Systems). The lysate and supernatant were processed and obtained similar to the RPA14-32 purification protocol, and the supernatant was incubated with Anti-FLAG beads for 2 h. The beads were washed with Wash Buffer (20 mM HEPES, 300 mM NaCl, 0.01% NP-40, 1 mM DTT) and eluted with Elution Buffer [20 mM HEPES, 150 mM NaCl, 0.01% NP-40, 25 μM EDTA, 1 mM DTT, 0.5 mg $mL^{-1}$ 3x FLAG peptides (APExBIO)]. The elute was loaded to Superdex 200 column (Cytiva) using the SEC buffer. The protein peaks were analyzed using SDS-PAGE gel and major protein fractions were concentrated. The concentrated fractions were transferred to Wash Buffer having 25 mM NaCl and loaded into the HiTrap Q HP column (Cytiva). RPA70 was eluted using a gradient of 25 mM to 1 M NaCl. The elution peak fractions were analyzed using SDS-PAGE. Fractions with adequate purity were pooled together and concentrated.

All the abovementioned protein purifications were quantified by measuring the sample absorbance at 280 nm using a Nanodrop spectrophotometer (ThermoFisher Scientific). The elutes were exchanged into similar buffer but containing 10% glycerol before snap-frozen in small aliquots for storage in –80 °C.

### Expression and purification of recombinant human TPP1-POT1-TIN2 (TPT) complex

The recombinant human TPP1-POT1-TIN2 (TPT) complex and TPP1 OB-domain protein were expressed using insect and bacterial cells, respectively. We followed the flashBAC ULTRA recommended protocol to generate a single baculovirus encoding all three subunits of TPT. The P3 baculovirus is titered by flow cytometric analysis for gp64 expression (Expression Systems). Typically, 1-2 L of *T. ni* cells (Expression Systems) were infected with the baculovirus at a cell density of 2 x $10^6$ cell $mL^{-1}$ and at a multiplicity of infection (MOI) of approximately 2-3. Cells were harvested for TPT purification 66-68 h post-infection. We purified the TPT complex following a previously reported protocol for human shelterin complexes (3). The purified TPT protein concentration is measured using absorbance at 280 nm before dispensing the proteins into 5 μL aliquots and snap-freezing using $LN_2$ for storage.

### AlphaFold2 prediction

AlphaFold2 predictions were computed using ColabFold (4) [version 1.5.2 from SBGrid (5)] on a local GPU server. The sequences used for computation are NP_937983.2 (TERT), NP_002936.1 (RPA70), NP_002937.1 (RPA32), NP_002938.1 (RPA14), and NP_001075955.2 (aa 1-197, $TPP1_{OB}$). The computation parameters used were --model-type auto --amber --num-relax 5 --use-gpu-relax.

### Molecular dynamics simulation for AlphaFold2 model relaxation

The starting structure for the MD simulations were taken from the PDB structure 7TRE (6) with residues 1-178 of TERT (TEN domain) and RPA32 domain modeled from the AlphaFold2 prediction. The DNA chain of 7TRE is excluded in the model. ACE and NME caps were added to the residues where the protein chains broke in the experimental structure. The protonation states of the residues were determined by PROPKA-3.5.1 (7, 8). The MD simulations were run using GROMACS 2022.5 (https://doi.org/10.5281/zenodo.7586780) using the CHARMM36 force field (updated July 2022) (9). The TIP3P water model was used as the solvent with sodium and chloride used to neutralize the system and with a 0.15 M concentration using a box size with 1 nm from the solute to the box edge (10, 11). The system had a total of 280,816 atoms with 84,381 waters, 413 sodiums, and 258 chlorides. The neighbor list, Lennard-Jones interactions, and Coulomb interactions were all cut off 1 nm and the long-range interactions were computed with particle-mesh Ewald (PME) (12). The system was energy minimized with 50,000 steps using the steepest descent algorithm. After, the system was equilibrated for 5 ns in the NVT ensemble with position restraints on the protein and nucleotide heavy atoms followed by 5 ns of NPT equilibration with the same position restraints using a force constant of 1,000 kJ/mol. Then 5 trajectories of 200 ns MD production runs were performed for each system with no position restraints under NPT conditions. The leap-frog integrator was used for equilibration and production using a 2 fs timestep using LINCS to constraint the hydrogen atom bonds (13). The Velocity rescale thermostat (14) was used to maintain the simulation at 300 K with a 0.1 ps time constant. The pressure was set to 1 bar which was maintained with the Parrinello-Rahman barostat (15) using a time constant of 2 ps and isotropic coupling.

### Direct telomerase primer extension assay

A standard base reaction (20 µl) consisted of dNTPs, 200 nM 5'Phos-TTAGGGTTAGGGTTAGGG DNA primer (Integrated DNA Technologies), and approximately 0.5 nM telomerase (quantified using TR dot blot) in the assay buffer containing 50 mM of HEPES–NaOH (pH 7.5), 100 mM KCl, 5 mM $MgCl_2$, and 2 mM DTT. Recombinant protein complexes and proteins were added to the reactions at concentrations specified in each figure. The reaction mixtures were pre-incubated at room temperature for 30 min before the enzymatic reactions were initiated by adding dNTPs. The dNTP mixture used was: 10 µM dGTP, 10 µM dTTP, 2.42 µM dATP, and 0.084 µM [α-$^{32}$P]dATP. The reactions were incubated at 37 °C for 1 h before they were quenched by 5 M ammonium acetate and 200 µg $mL^{-1}$ glycogen (Roche) for DNA precipitation. A radiolabeled 15-nt (TTAGGGTTAGGGTTA) oligonucleotide was added as a loading control in this step. After DNA precipitation, each sample was dissolved in 5 µl ultrapure water (Invitrogen #10977015) and 5 µl 2x formamide loading dye. Nine µl of each sample was loaded into each well of a 10 or 12% 1x TBE 7 M urea PAGE gel. Electrophoresis was performed at a constant 45 W until the bromophenol blue dye reached a third of the way from the gel bottom. The gels were vacuum-dried at 80 °C for 1 h before exposure to a storage phosphor screen. After exposure, the gels were imaged on an Amersham Typhoon gel imager (Cytiva). Enzyme activity and repeat addition processivity analysis was carried out using the GelAnalyzer software (version 23.11, www.gelanalyzer.com).

A standard base reaction (20 µl) consisted of dNTPs, 200 nM 5'Phos-TTAGGGTTAGGGTTAGGG DNA primer (Integrated DNA Technologies), and approximately 0.5 nM telomerase (quantified using TR dot blot) in the assay buffer containing 50 mM of HEPES–NaOH (pH 7.5), 100 mM KCl, 5 mM $MgCl_2$, and 2 mM DTT. Recombinant protein complexes and proteins were added to the reactions at concentrations specified in each figure. The reactions were pre-incubated at room temperature for 30 minutes. The enzymatic reactions were then initiated by adding dNTPs. The dNTP mixture used was: 10 µM dGTP, 10 µM dTTP, 2.42 µM dATP, and 0.084 µM [α-32P]dATP. The reactions were incubated at 37 °C for 1 h before they were quenched by DNA precipitation. A radiolabeled 15-nt (TTAGGGTTAGGGTTA) oligonucleotide was added as a loading control in this step. After DNA precipitation, each sample was dissolved in 5 µl ultrapure water (Invitrogen #10977015) and 5 µl 2× formamide loading dye and 9 µl of it was loaded into each well of a 10% 1× TBE 7 M urea PAGE gel. Electrophoresis was performed at a constant 45 W until the bromophenol blue dye reached a third of the way from the gel bottom. The gels were vacuum-dried at 80 °C for 1h before exposure to a storage phosphor screen. After exposure, the gels were imaged on a Amersham Typhoon gel imager (Cytiva). Enzyme activity analysis was carried out using the GelAnalyzer software (version 23.11, www.gelanalyzer.com).

### Fluorescence polarization DNA-binding assay

The 3xTEL DNA oligo was synthesized with a 5' Alexa-488 fluorescent dye (5'-Alexa488N-TTAGGGTTAGGGTTAGGG) (Integrated DNA Technologies). Each 40 µL reaction contained 1 nM fluorescent oligo in a buffer consisting of 50 mM HEPES (pH 7.5), 100 mM KCl, 5 mM $MgCl_2$, and 1 mM TCEP. The substrate was incubated with protein (concentrations as stated in related figures) for 1 h at 25 °C before fluorescence polarization measurement in a 384-well plate (Corning) using a Tecan Infinite M1000Pro plate reader. Fluorescence was measured with excitation at 485 nm and emission at 535 nm. Dissociation constants were determined from curve fitting using the Specific binding with Hill slope equation in GraphPad Prism 10 analysis software (version 10).

### Western blot analysis

To analyze phosphorylation of endogenous RPA, plasmid with 3xFLAG-Twin-Strep-TERT along with TR-HDV plasmid in a 1:3 ratio was transiently transfected in HeLa-EM2 cells (16) using Lipofectamine 2000 (Invitrogen). Forty-four hours later, control cells were treated with 50 µM etoposide and cultured further for 4 h. Cells were harvested, washed with 1xPBS and lysed in ice-cold RIPA buffer containing 50 mM Tris-HCl (pH 7.5), 150 mM sodium chloride, 1% NP-40, 1% sodium deoxycholate, 0.1% SDS, 1 mM DTT, supplemented with complete protease inhibitor cocktail (Roche). Cell lysates were incubated on ice for 45 min and clarified by centrifuging at 14,000 x g for 10 min at 4 °C. The clarified lysates were mixed with 4x NuPAGE LDS sample buffer (Invitrogen) supplemented with 50mM DTT and boiled at 95°C for 5 min before resolving them on SDS-PAGE (Mini-PROTEAN TGX 4–20% gels, BIO-RAD). Proteins were transferred onto nitrocellulose membranes, and the membranes were blocked with StartingBlock™ (TBS) blocking buffer (ThermoFisher Scientific) solution for 1 h at room temperature. The membranes were incubated with antibodies for FLAG M2-peroxidase mouse monoclonal antibody (Sigma Aldrich; A8592) and RPA32 rat monoclonal antibody (Cell signaling; 2208) diluted in StartingBlock™ (TBS) blocking buffer overnight at 4 °C. Horseradish peroxidase-conjugated secondary α-mouse (Santa Cruz Biotechnology; sc-525409), or α-rat antibodies (Cell signaling; 7077), and SuperSignal™ West Pico PLUS chemiluminescence substrate (Thermo Scientific) were used for chemiluminescence detection. For β-actin detection, the nitrocellulose membrane was stripped using Restore™ PLUS Western Blot Stripping Buffer (Thermo Scientific) for 15 min at room temperature and washed twice with PBS supplemented with 0.05% Tween 20 (Sigma). Following blocking for an hour at room temperature with StartingBlock™ (TBS) blocking buffer the membrane was incubated with β-actin mouse antibody (biotechne; MAB8929). The further chemiluminescence detection was carried out as stated above.

**Immunofluorescence (IF) and IF-fluorescence in-situ hybridization (IF-FISH) assays**

HeLa-EM2 cells for microscopy-based experiments were cultured on 8-well chambered glass bottom dishes (Cellvis, C8-1.5H-N). For RPA IF analysis, 3xFLAG-Twin-Strep-TERT or mutants with TR-HDV plasmids in a 1:3 molar ratio were transiently transfected in the HeLa-EM2 cells using Lipofectamine 2000 (Invitrogen). Cells were harvested 48 h later and pre-extracted with ice-cold Triton X-100 buffer containing 0.5% Triton X-100 (Sigma-Aldrich), 20 mM HEPES (pH 8), 50 mM NaCl, 3 mM $MgCl_2$, 300 mM sucrose for 4 min on ice before fixing them with 4% buffered formaldehyde solution at room temperature for 10 min. The cells were washed thrice with tris-buffered saline (TBS) and incubated overnight at 4 °C in blocking solution in PBS containing 3% goat serum, 0.2% triton X-100. Cells were incubated with primary antibodies (1:500) against FLAG tag (Sigma Aldrich; F1804), RPA32 (Cell signaling; 2208), and RAP1 (Novus Biologicals; NB100-292) or Coilin (Cell signaling; 14168) in blocking solution for 1h at room temperature. Glass dishes were washed thrice with PBS containing 0.2% triton X-100 and fluorophore-conjugated secondary-antibodies (Fisher Scientific; PIA32723, ThermoFisher Scientific; A-21245 and A-11035) (1:500) were applied for 1 h at room temperature. Glass dishes were subsequently washed thrice with PBS containing 0.2% triton X-100 with the first wash supplemented with 0.2 $\mu g\ ml^{-1}$ HOECHST stain. The fixed cells were maintained in PBS for imaging. The mounted glass dishes were imaged using an inverted Nikon fluorescence TIRF microscope equipped with a Kinetix sCMOS camera (Photometrics) and 100X oil-immersion objective. The IF image acquisition was performed on NIS-Elements (Nikon) and processed with ImageJ (17) with colocalizations quantified using CellProfiler (18). We used n = 80-110 nuclei for each sample as similar numbers were sufficient in previous studies to reveal phenotypic differences.

For IF experiments with RPA inhibitor TDRL-505 (Sigma-Aldrich; 530535), 3xFLAG-Twin-Strep-TERT or 3xFLAG-Strep-$TERT_{RTT}$ with TR-HDV plasmids in a 1:3 ratio were transiently transfected in HeLa-EM2 cells using Lipofectamine 2000 (Invitrogen). Forty-two hours later, cells were treated either with 100 mM TDRL-505 or DMSO and cultured further for 6 h. IF sample processing and data analysis was carried out as stated above.

For 53BP1 IF analysis, 3xFLAG-Twin-Strep-TERT and TR-HDV plasmids in a 1:3 ratio were transfected in HeLa-EM2 cells using Lipofectamine 2000 (Invitrogen). Forty-four hours later, control cells were treated with 50 mM etoposide and cultured further for 4 h. Cells were harvested and fixed with 4% buffered formaldehyde solution at room temperature for 10 min. The cells were washed thrice with tris-buffered saline (TBS) and incubated overnight at 4 °C in blocking solution in PBS containing 3% goat serum, 0.2% triton X-100. Cells were incubated with primary antibodies against 3xFLAG tag, RPA32, and 53BP1 (Novus Biologicals; NB100-304) in blocking solution for 1 h at room temperature. Glass dishes were washed thrice with PBS containing 0.2% triton X-100 and fluorophore-conjugated secondary-antibodies were applied for 1 h at room temperature. The cells were fixed again with 4% buffered formaldehyde solution at room temperature for 10 min and dehydrated with 70%, 95%, and 100% ethanol for two min each consecutively. Denaturation was carried out by adding hybridizing solution containing PNA probe (1:1,000) at 80 °C for 10 min. Glass dishes were subsequently washed thrice with PBS containing 0.2% triton X-100 with the first wash supplemented with 0.2 μg $ml^{-1}$ HOECHST stain. The fixed cells were maintained in PBS for imaging. The image acquisition and data analysis were done similarly as stated above.

### **CRISPR/Cas9-mediated TERT insertion at the *AAVS1* locus**

HEK293T cells (ATCC) were maintained at 37 °C under 5% $CO_2$ in Dulbecco's Modified Eagle Medium (DMEM) supplemented with 10% (v/v) fetal bovine serum, 100 μg $mL^{-1}$ penicillin/streptomycin, and 2 mM GlutaMAX (Invitrogen).

Our genome editing strategy is based on a previously reported protocol (19). The AAVS1 targeting gRNA sequence (T2 sequence: GGGGCCACTAGGGACAGGAT) was inserted into a mammalian expression vector that also encodes the Cas9 nuclease (Addgene, #48138). The Cas9 sequence has a C-terminal 2A-GFP sequence, which enabled us to screen for transfected cells using GFP detection. The donor plasmids were made by inserting TERT (WT and mutants) sequences by Gibson Assembly into the open reading frame (ORF) of a donor plasmid that harbors AAVS1 flanking sequences (Addgene, #68375). The ORF has an N-terminal 3xFLAG-Twin-Strep tag (making 3xFLAG-Twin-Strep-TERT) for western blot detection. There is also a puromycin resistance gene for selection of edited cells.

The Cas9/gRNA and donor plasmids were co-transfected into HEK293T cells, in a 6-well format, using the Jetprime DNA transfection reagent (Polyplus). At 48-72 h post-transfection, edited cells were selected with 1 μg $mL^{-1}$ puromycin. After two weeks, the resistant cells were tested for edited genome and 3xFLAG-Twin-Strep-TERT expression by PCR genotyping and western blot analysis, respectively. The genotyping used "inside-out" primer pairs (dna803-puromycin and dna804-puromycin) to detect DNA insertion into the targeted genomic loci. The primer sequences are:

dna803: TCGACTTCCCCTCTTCCGATG

dna804: GGTGCTAGGGCCGGGATTCTC

puromycin: GGTGCTAGGGCCGGGATTCTC

FLAG antibody (M2, Sigma-Aldrich) was used to detect exogenous expression of 3xFLAG-Twin-Strep-TERT in the cell lysates that were prepared using CHAPS lysis buffer. The validated edited cells were passaged in DMEM supplemented with 1 μg $mL^{-1}$ puromycin.

### **Telomere length Southern blot assay**

Genomic DNA was extracted from human cultured cells using a cell culture genomic DNA extraction kit (Omega Bio-tek) and quantified using a Qubit fluorometer (ThermoFisher Scientific). The digested genomic DNA samples were separated by pulse-field gel electrophoresis (Pippin Pulse, Sage Science). A high-molecular weight DNA marker (1 kb Extend DNA ladder, New England Biolabs) was labeled with Digoxigenin using a labeling kit (#MIR-3300, Mirus Bio) and purified by ethanol precipitation. Approximately 500 ng of DNA samples and 25 ng of marker were loaded into each well of the gel, and electrophoresis was performed for 9 hours using the pre-set 0.5-15kb protocol. Following electrophoresis, we proceeded with the Southern blot assay according to an established protocol (20). For hybridization, we used a $(TTAGGG)_6$ DNA oligo probe, labeled with digoxygenin-dUTP using terminal transferase (New England Biolabs). The telomeric probe was incubated with the blot at a concentration of 1 nM for 1 hour at 42 °C. Anti-DIG Fab-Alkaline Phosphatase was used to immunoblot the hybridized telomeric probe. After several washes, CDP-Star (Sigma-Aldrich) was added for chemiluminescent detection. Imaging was performed using a cooled CCD camera (Azure C300, Azure Biosystems) with a typical exposure time of approximately 30 minutes.

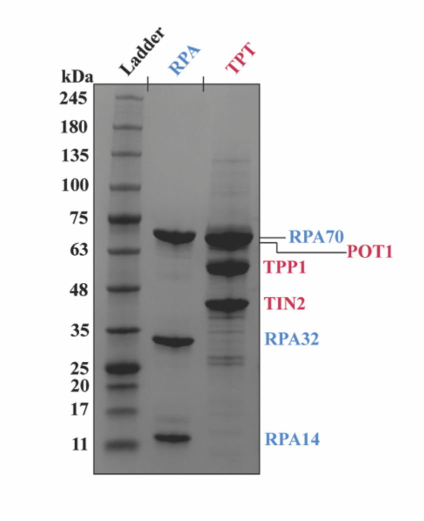


**Fig. S1. SDS-PAGE analysis of recombinant human RPA and TPP1-POT1-TIN2 proteins.** SDS-PAGE analysis of purified recombinant human RPA and TPP1-POT1-TIN2 (TPT) protein complexes. RPA and TPT subunits are colored in blue and red respectively.

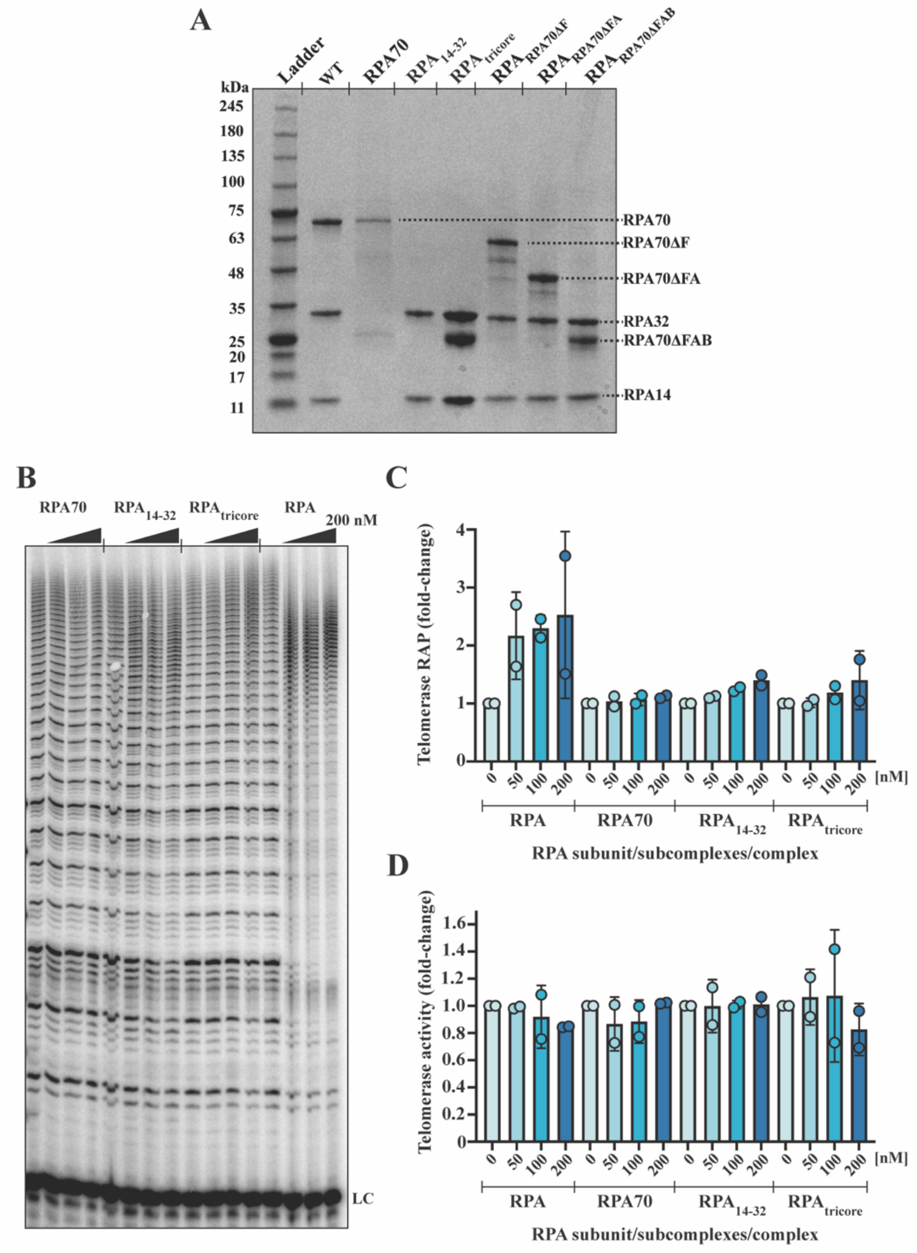


**Fig. S2. Effect of RPA truncation mutants and subcomplexes on telomerase processivity.** **(A)** SDS-PAGE analysis of purified recombinant RPA subunit, truncated complexes and subcomplexes. $RPA_{tricore}$ complex contains RPA70 OB-C domain, RPA32 and RPA14 subunits. **(B)** Telomerase primer extension assay with 50, 100, 200 nM addition of RPA70, $RPA_{14\text{-}32}$, $RPA_{tricore}$, and RPA (full-length control). DNA oligo, $(TTAGGG)_3$, was used as the primer for the primer extension assay. Hot dATP was used to label telomerase extended primers. LC: loading control. **(C)** Quantification of telomerase repeat addition processivity, which is calculated as the ratio of ≥10 repeat products to <10 repeat products, based on gel band intensities. **(D)** Quantification of total telomerase activity, calculated by summing the intensities of all product bands on the gel.

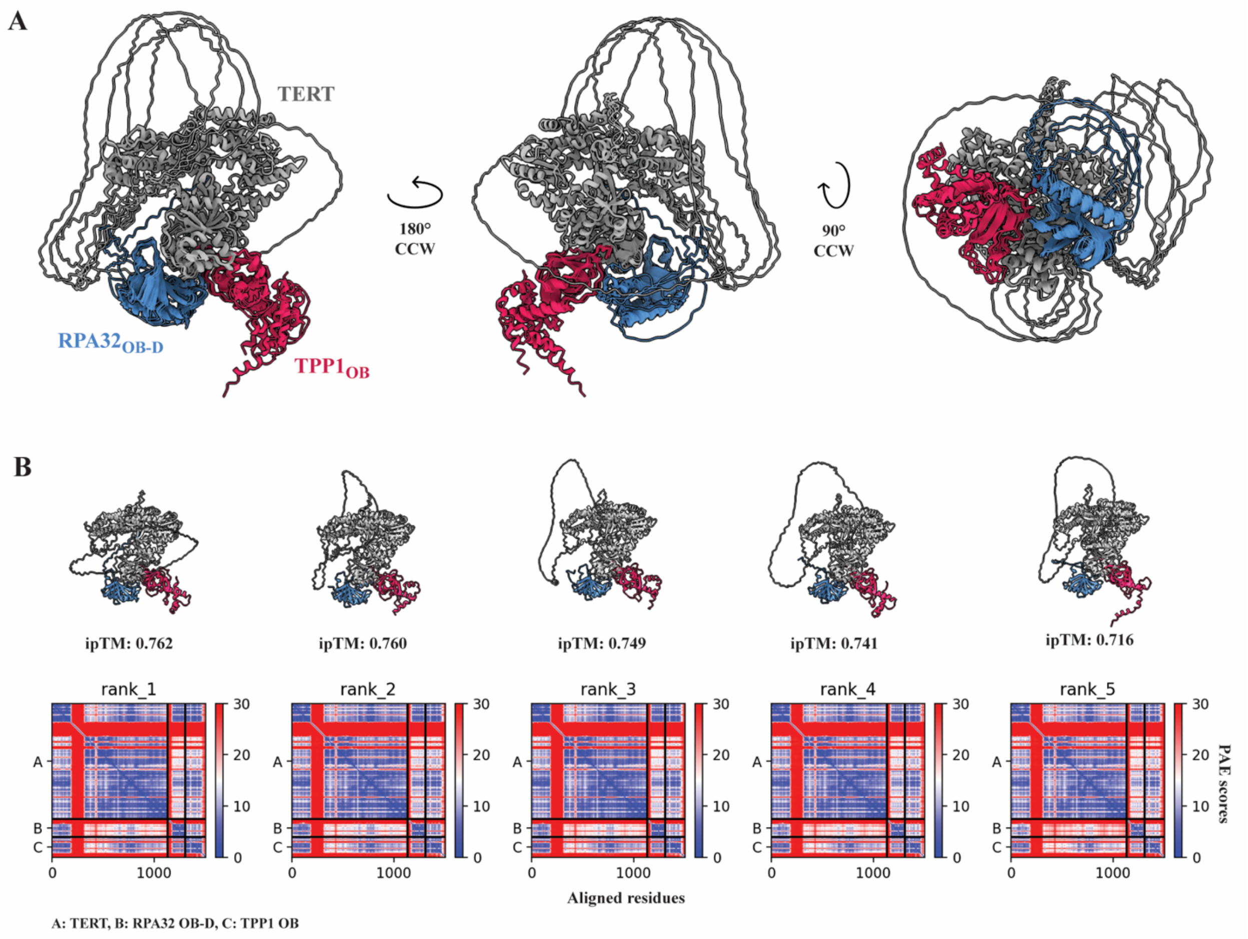


**Fig. S3. AlphaFold2 multimer prediction for TERT-RPA32$_{OB\text{-}D}$-TPP1$_{OB}$ co-complex. (A)** Aligned models from five AlphaFold2 multimer prediction of TERT-RPA$_{32OB\text{-}D}$-TPP1$_{OB}$ interactions. R.M.S.D between the top ranked model and the rest of the models ranges from 0.5 to 0.8 Ångstroms (Å). CCW: counterclockwise. **(B)** The individual predicted models with their corresponding integrated predicted TM-score (ipTM) and Predicted Aligned Error (PAE) plot. Unit for the PAE plots (heatmaps) is Å.

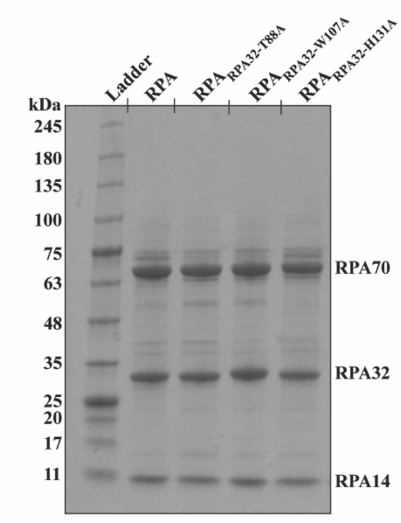


**Fig. S4. SDS-PAGE analysis of recombinant human RPA mutants.** SDS-PAGE analysis of purified recombinant human RPA containing RPA32 mutations identified from AlphaFold2 TERT-RPA32 multimer predictions.

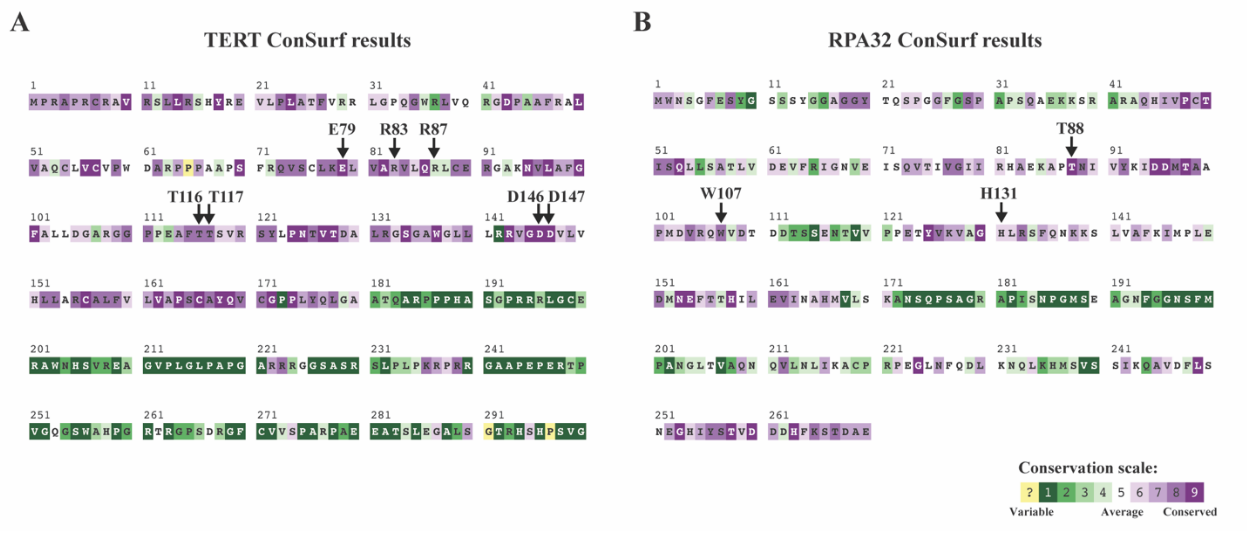


**Fig. S5. Sequence conservation analysis for TERT and RPA32.** ConSurf analysis for **(A)** TERT and **(B)** RPA32 sequences. Only the relevant region of analysis is shown for TERT. Residues tested in this work are marked as down arrows.

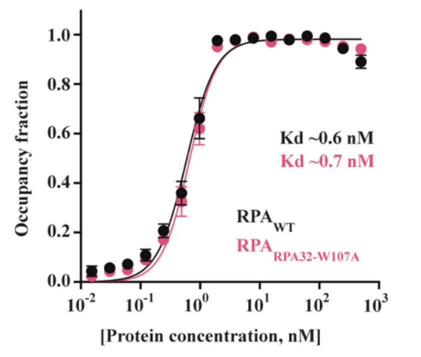


**Fig. S6. DNA-binding assay for characterizing WT and mutant RPA binding to telomeric single-stranded DNA.** Fluorescence polarization binding assay was used to quantify the binding affinity of WT RPA and mutant RPA (containing W107A mutation in the RPA32 subunit) to a Alexa488-labeled 3xTEL (TTAGGGTTAGGGTTAGGG) oligo. The binding affinities, Kd, were calculated by fitting the datapoints to a single-binding site equation.

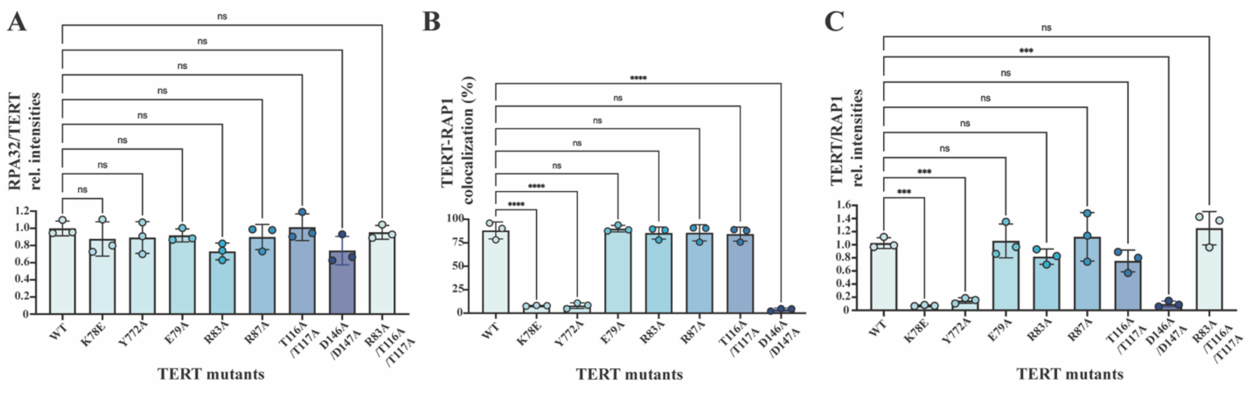


**Fig. S7. Quantification of RPA32 to TERT and TERT to RAP1 colocalization in immunofluorescence imaging experiments. (A)** Relative intensity analysis of RPA32/TERT fluorescence puncta. **(B)** Colocalization analysis of TERT on RAP1 puncta. RAP1 is used to mark telomeres in nuclei. **(C)** Relative intensity analysis of TERT/RAP1 fluorescence puncta. ns stands for nonsignificant; $P > 0.05$, *** stands for $P \leq 0.001$, and **** stands for $P \leq 0.0001$.

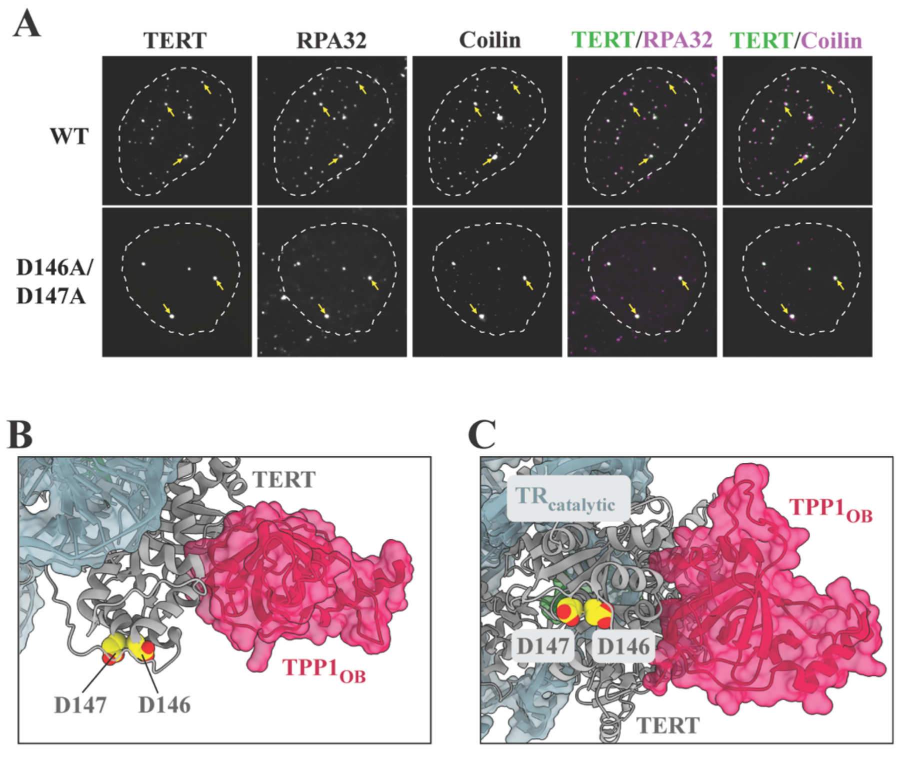


**Fig. S8. Cajal bodies colocalization and structural analysis of TERT D146A/D147A mutant.** **(A)** Immunofluorescence image analysis of WT and D146A/D147A mutant TERT colocalization with neo-Cajal and Cajal bodies in HeLa cells. WT TERT, detected by FLAG antibodies, colocalized with neo-Cajal bodies (inferred from >12 puncta), which were marked by Coilin antibodies. In contrast, the D146A/D147A mutant localized to Cajal bodies (inferred from 2-4 puncta per nucleus). RPA32 colocalized with TERT for WT and D146A/D147A at these Cajal bodies. Representative TERT puncta are marked by yellow arrows and mapped onto individual fluorescence channels and merged images. The nuclei boundaries are demarcated by white dashed lines. (**B & C**) TERT D146 and D147 are highlighted as yellow- and heteroatom-colored space-filled residues in the Telomerase-TPP1$_{OB}$ cryo-EM structure (PDB: 7TRE) to illustrate the spatial distance between the two sites.

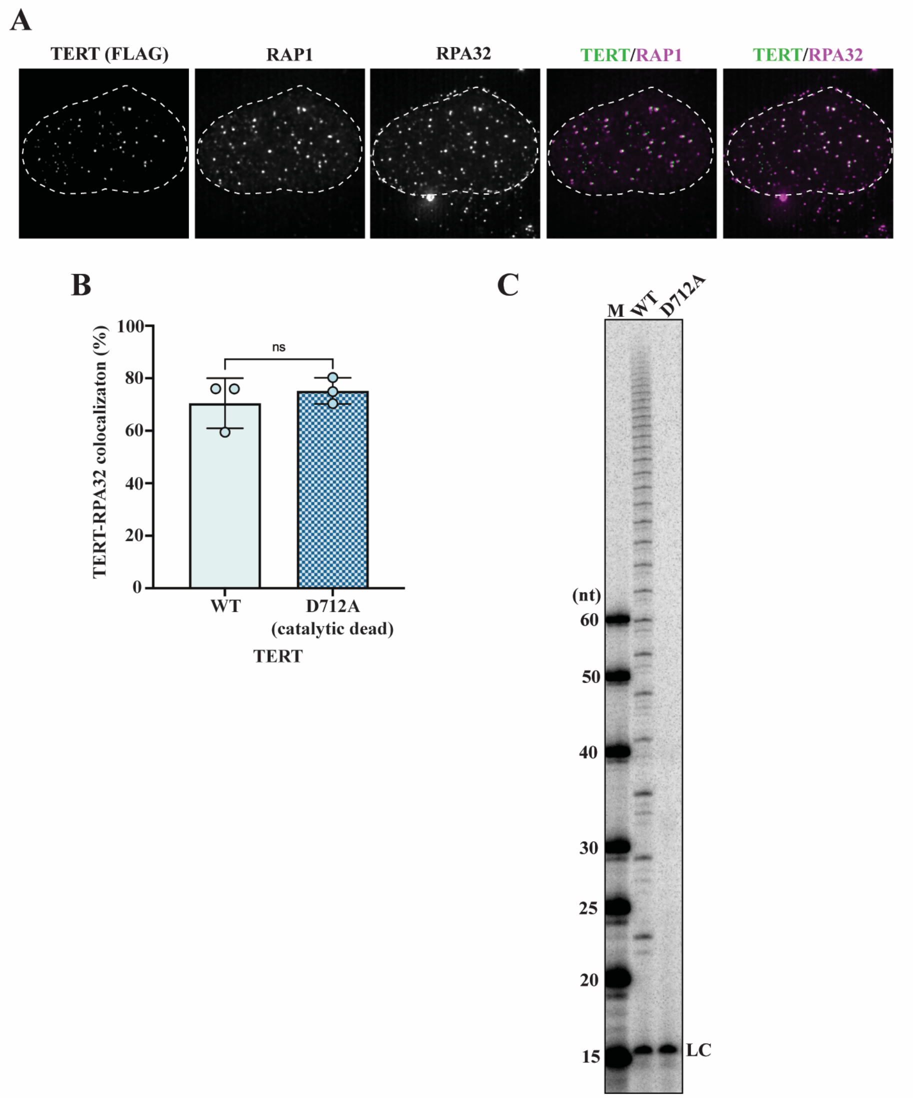


**Fig. S9. RPA-telomere colocalization analysis of cells expressing catalytically dead telomerase. (A)** Immunofluorescence colocalization image analysis of the catalytically dead D712A TERT mutant in HeLa cells. RPA32 colocalized with TERT at telomeres, which are marked by RAP1 antibodies. The nucleus boundary is demarcated by white dashed lines. (**B**) Quantification of TERT and RPA32 colocalization at telomeres. **(C)** Direct telomerase assay comparing activity of WT and D712A TERT mutant. M: marker and LC: loading control. ns stands for nonsignificant.

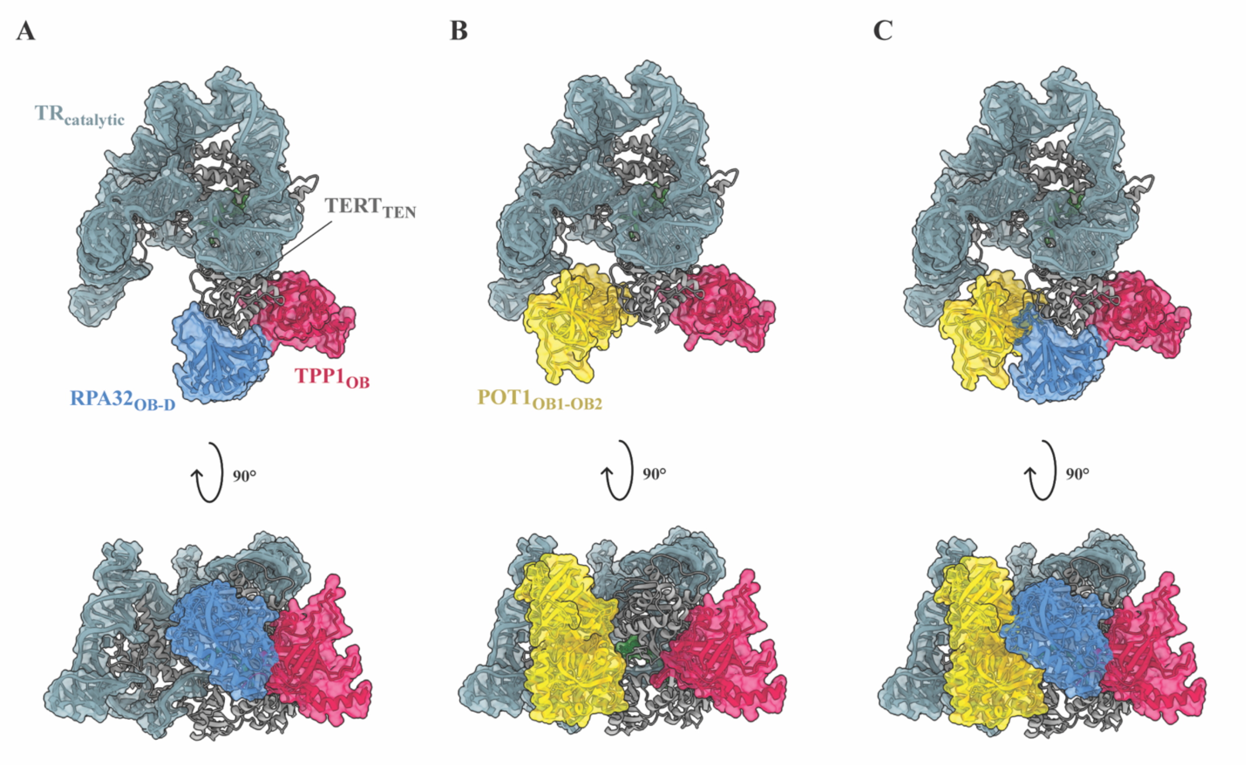


**Fig. S10. Structural compatibility analysis of RPA32 and POT1 binding to TERT. (A)** The AlphaFold2 predicted TERT-RPA32$_{OB-D}$ model mapped onto TERT-TR-TPP1$_{OB}$ cryo-EM structure (PDB: 7TRE). (**B**) A cryo-EM structure of TERT-TR-TPP1$_{OB}$-POT1$_{OB1-OB2}$ (PDB: 7QXS) oriented the same as panel **A**. **(C)** The AlphaFold2 RPA32$_{OB-D}$ model mapped onto the TERT-TR-TPP1$_{OB}$-POT1$_{OB1-OB2}$ cryo-EM structure.

| Oligo Name | Oligo Sequence (5'-3') |
|---|---|
| **Y772A_circ_F** | CCTCCAGCCGgcgATGCGACAGTTC |
| **Y772A_circ_R** | TCTGTCAAGGTAGAGACG |
| **K78E_circ_F** | GTCCTGCCTGgaaGAGCTGGTGG |
| **K78E_circ_R** | ACCTGGCGGAAGGAGGGG |
| **TT_circ_F** | gcgAGCGTGCGCAGCTACCTGCCCAACA |
| **TT_circ_R** | ggcGAAGGCCTCGGGGGGGCCCCCGCG |
| **DD_circ_R** | gcgGTGCTGGTTCACCTGCTGGCACGC |
| **DD_circ_F** | ggcGCCCACGCGGCGCAGCAGCAGCCCCC |
| **T116I_circ_F** | accAGCGTGCGCAGCTACCTGCCCAACA |
| **T116I_circ_R** | gatGAAGGCCTCGGGGGGGCCCCCGCG |
| **F115L_circ_F** | ctgACCACCAGCGTGCGCAGCTACCTG |
| **F115L_circ_R** | GGCCTCGGGGGGGCCCCCGCG |
| **D712A_circ_F** | gccGTGACGGGCGCGTACGACACCATCCCC |
| **D712A_circ_R** | CACCTTGACAAAGTACAGCTCAGGCGG |
| **E79AInsF** | AGGTGTCCTGCCTGAAGGCGCTGGTGGCCCGAG |
| **E79AInsR** | TCAGTCCAGGATGGTCTTGAAGTCTGA |
| **E79AvecF** | CTTCAGGCAGGACACCT |
| **E79AvecR** | TCAGACTTCAAGACCATCCTGGACTGA |
| **R83AInsF** | CTGAAGGAGCTGGTGGCCgcgGTGCTGCAGAGGCTGTGCGA |
| **R83AVecF** | GGCCACCAGCTCCTTCAG |
| **R83P_Ins_F** | CTGAAGGAGCTGGTGGCCccaGTGCTGCAGAGGCTGTGCGA |
| **R87A_circ_F** | AGTGCTGCAGgcgCTGTGCGAGCG |
| **R87A_Circ_R** | CGGGCCACCAGCTCCTTC |
| **RPA2_T88A_circR** | tggagccttctctgcatgtctgatgatccc |
| **RPA2_T88A_circF** | gccaacattgtttacaaaatagatgacatgacagct |
| **RPA2_W107A_circR** | ctggcgaacgtccatgggtgcagc |
| **RPA2_W107A_circF** | gccgttgacacagatgacaccagcagtg |
| **RPA2_H131A_circR** | gcctgccactttcacatatgtttctggaggaac |
| **RPA2_H131A_circF** | gccctgagatcttttcagaacaaaaagagcctggtagcc |
| **hTR_NBprobe1** | gactcgctccgttcctcttc |
| **hTR_NBprobe2** | gctctagaatgaacggtggaa |
| **hTR_NBprobe3** | cctgaaaggcctgaacctc |
| **hTR_NBprobe4** | cgcctacgcccttctcagt |
| **hTR_NBprobe5** | atgtgtgagccgagtcctg |
| **RPA1_delOB_R** | gagactccctccggacttgtc |
| **RPA1_delOBF_F** | ggactcgggcagccgcaagtagc |
| **RPA1_delOBFAB_F** | gggagtaacaccaactggaaaaccttg |
| **RPA1_delOBFA_F** | gctaacaagcagttcacagctgtt |

**Table S1. Primer DNA sequences for mutagenesis**

### References


65. G. Cristofari, J. Lingner, Telomere length homeostasis requires that telomerase levels are limiting. *EMBO Journal* **25** (2006).

66. L. A. Henricksen, C. B. Umbricht, M. S. Wold, Recombinant replication protein A: Expression, complex formation, and functional characterization. *Journal of Biological Chemistry* **269** (1994).

67. C. J. Lim, A. J. Zaug, H. J. Kim, T. R. Cech, Reconstitution of human shelterin complexes reveals unexpected stoichiometry and dual pathways to enhance telomerase processivity. *Nat Commun* **8** (2017).

68. M. Mirdita, K. Schütze, Y. Moriwaki, L. Heo, S. Ovchinnikov, M. Steinegger, ColabFold: making protein folding accessible to all. *Nat Methods* **19** (2022).

69. A. Morin, B. Eisenbraun, J. Key, P. C. Sanschagrin, M. A. Timony, M. Ottaviano, P. Sliz, Collaboration gets the most out of software. *Elife* **2013** (2013).

70. B. Liu, Y. He, Y. Wang, H. Song, Z. H. Zhou, J. Feigon, Structure of active human telomerase with telomere shelterin protein TPP1. *Nature* **604** (2022).

71. C. R. Søndergaard, M. H. M. Olsson, M. Rostkowski, J. H. Jensen, Improved treatment of ligands and coupling effects in empirical calculation and rationalization of p K a values. *J Chem Theory Comput* **7** (2011).

72. M. H. M. Olsson, C. R. SØndergaard, M. Rostkowski, J. H. Jensen, PROPKA3: Consistent treatment of internal and surface residues in empirical p K a predictions. *J Chem Theory Comput* **7** (2011).

73. J. Huang, S. Rauscher, G. Nawrocki, T. Ran, M. Feig, B. L. De Groot, H. Grubmüller, A. D. MacKerell, CHARMM36m: An improved force field for folded and intrinsically disordered proteins. *Nat Methods* **14** (2016).

74. P. Mark, L. Nilsson, Structure and dynamics of the TIP3P, SPC, and SPC/E water models at 298 K. *Journal of Physical Chemistry A* **105** (2001).

75. W. L. Jorgensen, J. Chandrasekhar, J. D. Madura, R. W. Impey, M. L. Klein, Comparison of simple potential functions for simulating liquid water. *J Chem Phys* **79** (1983).

76. T. Darden, D. York, L. Pedersen, Particle mesh Ewald: An N·log(N) method for Ewald sums in large systems. *J Chem Phys* **98** (1993).

77. B. Hess, H. Bekker, H. J. C. Berendsen, J. G. E. M. Fraaije, LINCS: A Linear Constraint Solver for molecular simulations. *J Comput Chem* **18** (1997).

78. G. Bussi, D. Donadio, M. Parrinello, Canonical sampling through velocity rescaling. *Journal of Chemical Physics* **126** (2007).

79. M. Parrinello, A. Rahman, Polymorphic transitions in single crystals: A new molecular dynamics method. *J Appl Phys* **52** (1981).

80. I. Weidenfeld, M. Gossen, R. Löw, D. Kentner, S. Berger, D. Görlich, D. Bartsch, H. Bujard, K. Schönig, Inducible expression of coding and inhibitory RNAs from retargetable genomic loci. *Nucleic Acids Res* **37** (2009).

81. J. Schindelin, I. Arganda-Carreras, E. Frise, V. Kaynig, M. Longair, T. Pietzsch, S. Preibisch, C. Rueden, S. Saalfeld, B. Schmid, J. Y. Tinevez, D. J. White, V. Hartenstein, K. Eliceiri, P. Tomancak, A. Cardona, Fiji: An open-source platform for biological-image analysis. [Preprint] (2012). https://doi.org/10.1038/nmeth.2019.

82. A. E. Carpenter, T. R. Jones, M. R. Lamprecht, C. Clarke, I. H. Kang, O. Friman, D. A. Guertin, J. H. Chang, R. A. Lindquist, J. Moffat, P. Golland, D. M. Sabatini, CellProfiler: Image analysis software for identifying and quantifying cell phenotypes. *Genome Biol* **7** (2006).

83. M. Dalvai, J. Loehr, K. Jacquet, C. C. Huard, C. Roques, P. Herst, J. Côté, Y. Doyon, A Scalable Genome-Editing-Based Approach for Mapping Multiprotein Complexes in Human Cells. *Cell Rep* **13** (2015).

84. M. Kimura, R. C. Stone, S. C. Hunt, J. Skurnick, X. Lu, X. Cao, C. B. Harley, A. Aviv, Measurement of telomere length by the southern blot analysis of terminal restriction fragment lengths. *Nat Protoc* **5** (2010).
